# The central complex of the larval fruit fly brain

**DOI:** 10.1101/2025.09.30.679510

**Authors:** Laura Lungu, Nicolò Ceffa, Michael Clayton, Marta Zlatic, Albert Cardona

## Abstract

In holometabolous insects such as the fruit fly *Drosophila melanogaster*, the brain central complex (CX) develops during metamorphosis and serves the adult stage. Whether a form of the CX exists in the brain of the evolutionarily novel larval stages is not known. Here, we analyzed the connectome of the larval brain and, on the basis of neuronal lineages, synaptic connectivity patterns, and anatomy, identified a putative larval CX, comprising 4 key neuropils: the protocerebral bridge (PB), the ellipsoid body (EB), the fan-shaped body (FB) and the noduli (NO). Consistent with our interpretation, we found in the larval brain synaptic connectivity patterns characteristic of the adult, including (i) visual input into the PB and EB; (ii) modulation of CX neuropil inputs by the mushroom body (MB); (iii) reciprocal connectivity between CX neuropils and select MB compartments; and (iv) strong connectivity between CX neuropils. While some neuronal lineages contributing to the larval CX do not contribute to the adult CX, many others are conserved. The characterization of a larval CX brings structure to largely unexamined larval brain circuits, linking with a vast body of literature, and will inform the design of experiments to probe larval brain function.

## 1 Introduction

The Central Complex (CX) is a conserved set of neuropils found across arthropods that acts as a sensory integration area and a center for coordinated motor activity [1–3]. In *Drosophila melanogaster*, the functions of the CX include multisensory navigation, path integration, place learning, allocentric orientation of the head relative to its body, sleep regulation, and providing an internal sense of direction in the absence of stimuli, among others [1, 4–13].

The fruit fly larva exhibits a range of behaviors that require spatial navigation, including chemotaxis for foraging and escape [14–17] and aversive phototaxis in response to blue light [18–20]. In addition to responding to an isolated unimodal stimulus, the larva integrates competing stimuli prior to decision making for navigation [21]. Furthermore, larvae sleep, which is required for long-term memory [22] just like in the adult fly [23, 24], where sleep is regulated by the central complex [12].

Anatomically, the larval central brain shares the set of neuroblasts with the adult [25], and presents neuropil areas that directly correspond to those of the adult brain such as the Antennal Lobe (AL), the Mushroom Body (MB), and the Lateral Accessory Lobe (LAL), in addition to more larval-specific neuropils such as the larval optic neuropil (LON) [26]. The organization and synaptic connectivity of the larval AL, MB, and LON are well understood at present [27–29]. Nevertheless, these neuropils constitute only ∼30% of the complete larval brain connectome by number of neurons [30]. We postulate that amongst the remaining ∼70% of brain neurons, and given the reported conservation of neuroblasts [25] and neuronal identities from larval to adult [31], and the larval behaviors in navigation and sleep that in the adult are associated with the CX, the question of whether the larval brain harbors a simpler yet homologous set of Central Complex neuropils must be examined. Furthermore, in other holometabolous insect orders, neuroblast proliferation arrests at a later point in the stereotyped sequence of neuron types [32], generating more neurons embryonically and rendering the central complex identifiable at the larval stage, such as in the beetle *Tribolium castaneum* where the CX develops embryonically [33, 34]. In the fruit fly larva, hundreds of late embryonic-born neurons known to later on pioneer the development of the adult CX remain undifferentiated throughout larval life [35].

The adult fruit fly CX has four well-studied neuropils: the Protocerebral Bridge (PB), the Fan-shaped Body (FB), the Ellipsoid body (EB) and the Noduli (NO) [4] plus, as of recently, the Asymmetrical Body (AB) [36], in addition to several smaller accessory neuropils [8, 37, 38]. The adult Lateral Accessory Lobe (LAL) is a CX-associated neuropil interconnected with all its neuropils [38] and is therefore an interesting reference point since the larval brain also has an LAL [26].

The larval brain, though, does not present fused brain hemispheres like in the adult. With adult CX neuropils being largely medial structures, any putative larval brain counterparts will differ significantly by morphology alone. The basis of our search for the larval CX has to rely on (1) neuronal lineages generated by the same neuroblasts; (2) the relative spatial location of neurons within the overall neuropil since these are conserved for neuropils known to be homologous such as the AL, MB, LON and LAL; (3) the synaptic circuit architecture organizing the adult central complex neuropils; and (4) the patterns of sensory inputs. For (1) we use embryonic-born neurons that remain identifiable yet undifferentiated throughout larval life, a subset of which will develop into the adult CX [35]. For (2), (3) and (4) we rely on the fully reconstructed neuronal arbors constituent of the complete connectome of the larval brain [30], together with prior publications on larval sensory systems [27, 29, 39–44]. We use all four in an iterative process to progressively discover the complete list of neurons composing the putative larval CX.

## 2 Methods

### 2.1 Connectome data

The connectome of a 2-hour old first instar larval brain was used, as reconstructed previously by us [30]. The neuronal reconstructions, synapse labels, and neuron annotations, together with the electron microscopy volume, is available at the VirtualFlyBrain.

### 2.2 Fly Strains

The following effector lines were used: UAS-impTNT-HA, UAS-TNT-HA, obtained from the Bloomington Stock Center (Indiana, USA). We used the following split-GAL4 GMR driver lines: SS01978, SS26146, SS2856, SS22965, SS04497, SS25740, SS27197, SS36847 (HHMI Jannelia Research Campus; [45, 46]). See Fig. 10.

We used TH-GAL4, a line that tags all dopaminergic neurons in the central nervous system of the larva [47], and crossed it with tsh-GAL80 to suppress expression in the VNC. The following driver line was used: R57C10-FlpL in su(Hw)attP8;; pJFRC210-10XUAS-FRT>STOP>FRT-myr::smGFP-OLLAS in attP2. pJFRC210-10XUAS-FRT>STOP>FRT The following effector line was used: w; tsh-Gal80/cyo.Tb.RFP; TH-Gal4. These were crossed, and the desired progeny was: R57C10-FlpL / + ; tsh-Gal80 / CyO.Tb.RFP ; TH-Gal4 / UAS-smGFP. To select for larvae that express GAL4 only in the brain, only larvae with red fluorescent bodies were dissected, as this indicates the presence of tsh-GAL80.

### 2.3 Nomenclature

We classified neurons using the suffixes “.b” (axon boutons) and “.d” (dendrites) for CX neuropils. Hence, a neuron annotated as NO.b projects its axon into the NO, and likewise, a neuron annotated as FB.d places its dendrites in the FB. Neurons that are both .b and .d for CX neuropils we annotated as Core CX neurons.

### 2.4 Searching for adult brain synaptic connectivity patterns in the larval brain

We adopted a search strategy guided by circuit features of the adult brain CX neuropils. Since the adult brain presents fused hemispheres at the midline whereas the larva has not, we placed more emphasis of searching for circuit motifs rather than anatomical similarity, while preserving the relative spatial location of a CX neuropil as a constraint.

These included:

1. Integration of sensory inputs: we searched for synaptic pathways corresponding to the adult’s for integrating visual inputs into the PB and EB [38], and gustatory inputs into the FB.
2. Association between centers for memory and spatial navigation: we looked for the characteristic link between the MB, the center for associative memory, and CX neuropils, which includes synaspes from MBONs (MB output neurons) to the FB, EB and NO in the adult fly.
3. Recurrent intra-CX connectivity, such as with columnar neurons relating the PB with the FB and EB, or with the NO.

### 2.5 Identifying larval neuronal lineages that contribute to the larval CX

The *Drosophila melanogaster* central brain consists of stereotyped neural lineages, developmental-structural units of macrocircuitry formed by the sibling neurons of single progenitor cells, the neuroblasts [48], a subset of which structures the CX [49]. Starting with the currently identified set of quiescent, embryonic-born neurons present in the larval brain that develop into to the adult CX during metamorphosis [35] we identified larval brain neurons satisfying imposed connectivity rules as per known circuit patterns across the adult CX neuropils [8, 36–38].

We broadened the search beyond these lineages based on the expectation that some larval neurons may have been recruited to CX structures but would later take another identity in the adult, as is known for larval MB neurons [31], on the basis of matching synaptic connectivity patterns as observed in the adult; for example, for EB Ring neurons we looked for neurons with recurrent axo-axonic synapses with each other and with Wedge neurons, where their axon is located at the most anterior end of the brain neuropil, at an intermediate dorso-ventral position just anterior to the MB medial lobe.

### 2.6 Immunohistochemistry

For Immunohistochemistry, we adapted the protocol from HHMI Janelia Research Campus, in combination with the protocol from [50]. The following primary antibodies were used: **Mouse** *α***-Neuroglian; Rabbit** *α***-HA Tag; Rat** *α***-FLAG Tag** (Sigma). The following secondary antibodies were used: AF488 Donkey *α***-Mouse; DL549 Goat** *α***-Rabbit (Sigma)**.

The larval central nervous system (CNS) was dissected in cold 1× phosphate-buffered saline (PBS). The tissue was then transferred into 2 mL Protein LoBind tubes containing cold 4% paraformaldehyde (PFA) in 1× PBS, and incubated for 1 h at room temperature (RT) while nutating. The PFA was then removed and tissues were washed in 1.75 mL of 1% PBT (PBS with 0.3% Triton X-100) four times for 15 min each with nutation. Samples were then blocked with 5% Normal Donkey Serum (NDS Jackson Immuno Research; prepared as 95 *µ*L PBT + 5 *µ*L NDS), and incubated for 2 h at RT on a rotator with tubes upright. Primary antibody incubation was carried out in 1% PBT (typically 100 *µ*L per tube) for 4 h at RT, followed by two consecutive overnights at 4*^◦^*C with continuous rotation. After primary incubation, tissues were washed four times in 1.75 mL of 1% PBT for 15 min each.

Secondary antibody incubation was performed in 1% PBT (100 *µ*L per tube) for 4 h at RT, followed by 1–2 overnights at 4*^◦^*C with continuous rotation. Post-secondary washes were performed four times in 1.75 mL of 1% PBT for 15 min each. An additional blocking step with 5% Normal Mouse Serum (NMS; Jackson ImmunoResearch, #015-000-120) in PBT was carried out for 1.5 h at RT prior to overnight incubation with at 4*^◦^*C with the conjugated antibody. Following incubation, samples were washed four times in 1.75 mL of 1% PBT for 15 min each.

For mounting, tissues were placed on poly-L-lysine (PLL)-coated coverslips. Samples were dehydrated through a graded ethanol series (30%, 50%, 75%, 95%, and three changes of 100%), soaking for 10 min at each step. Tissues were cleared by immersion in three sequential 5 minute xylene baths. Finally, samples were embedded by applying 4-5 drops of dibutyl phthalate in xylene (DPX) to the mounted tissue, placing the coverslip (DPX side down) onto a prepared slide with spacers, and applying gentle pressure to seat the coverslip. Slides were left to dry in the hood for 1-2 days prior to imaging.

### 2.7 Behavioral Assays during visual stimulation

9 split GAL4 lines were used and crossed with either UAS-TNT (effector) or UAS-impTNT (control) genetic driver lines lines. The cross was set at 25*°*C and the flies were laid on fly food for 3 days. Larvae containing the UAS-TNT or UAS-impTNT transgene were raised at 18*°*C for 7 days with normal cornmeal food. Third instar larvae were used for all experiments.

Larvae were separated from food by using 15% sucrose and washed with water, then dried and placed in the center of the arena, consisting in 3% Bacto agar gel in a 25 × 25 cm square plastic plate. Experiments were conducted at 25*°*C. Larvae were monitored with the Multi-Worm Tracker (MWT) software (http://sourceforge.net/projects/mwt); [51].

For light-stimulation experiments, we used approximately 30 larvae for each run. The larvae were presented with green light for 40 seconds, and the amount of larvae turning was monitored before, during and after stimulus presentation. 6 runs were performed for every line.

### 2.8 Behavioral Quantification

Larvae were tracked in real-time using the MWT software [52]. We rejected objects that were tracked for less than 5 seconds or moved less than one larval body length. For each larva, MWT returns a contour, spine and center of mass as a function of time. From the MWT tracking data we computed the key parameters of larval motion, using Choreography software [52] with variables as in [51]. From the tracking data, we quantified crawling and rolling events and the speed of peristaltic crawling strides.

## 3 Results

### 3.1 Correspondence between larval and adult neuronal lineages

On the basis of neurons from lineages known to contribute to the adult central complex and the spatial arrangement of their arbors in the brain, we will now examine each larval central complex neuropil individually. For each CX neuropil we first summarize the main cell types and circuit characteristics in the adult, and then describe larval neurons that match them.

#### 3.1.1 The larval Protocerebral Bridge (PB)

In the adult *Drosophila*, the PB comprises two sets of bilaterally symmetric compartments, sometimes referred to as glomeruli 1–9 [38], positioned at the most posterior-dorsal location possible in the brain.

These compartments are arranged in a continuous manner medio-laterally, contacting at the midline. In the adult, about 600 neurons innervate the PB, organized into hundreds of types (194; [37]) that are split into two main general groups: the columnar neurons (from lineages DM1, DM2, DM3, DM4 and DM6) whose dendrites innervate one or more of the 9 + 9 compartments of the PB [37]; and the horizontal neurons (also known as horizontal fibers) derived from a single lineage (PBp1; [35]) whose axons innervate many or all PB compartments.

In the adult, the PB receives visual input via relay neurons (POL neurons) conveying information on polarized light, in a highly structured pattern across its compartments that binarizes the continuum of angles of polarized light ([54]). Then the PB relays this information to the EB compartments (Fig. 1).

**Figure 1:**
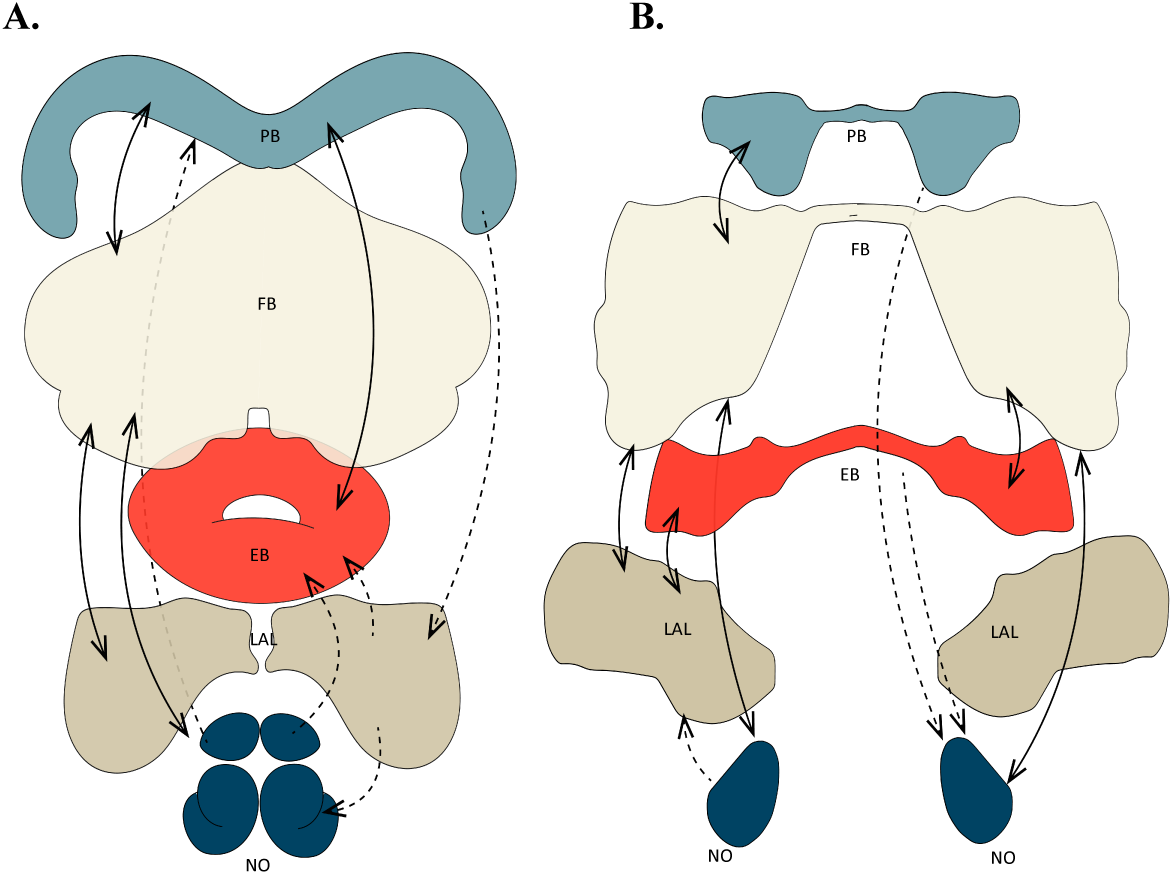
The Central Complex Neuropils in Adult and Larva. Dotted-line arrows represent unidirectional connections, filled-line arrows indicate bidirectional connections. In panel **A.** we show the adult *Drosophila* CX. Bidirectional connectivity is seen between PB-FB, PB-EB, FB-LAL and FB-NO, with unidirectional connections from NO-PB, PB-LAL, LAL-NO and NO-EB; Panel **B.** displays the larval *Drosophila* CX. Bidirectional connections are PB-FB, FB-EB, FB-LAL, FB-NO and EB-LAL; unidirectional connections are seen from NO-LAL, PB-NO and EB-NO.The volumes of the adult neuropils were adapted from [8]. The volumes of the larval neuropils were drawn via CATMAID using the .b and .d parts of the neurons contributing to each.

**Figure 2:**
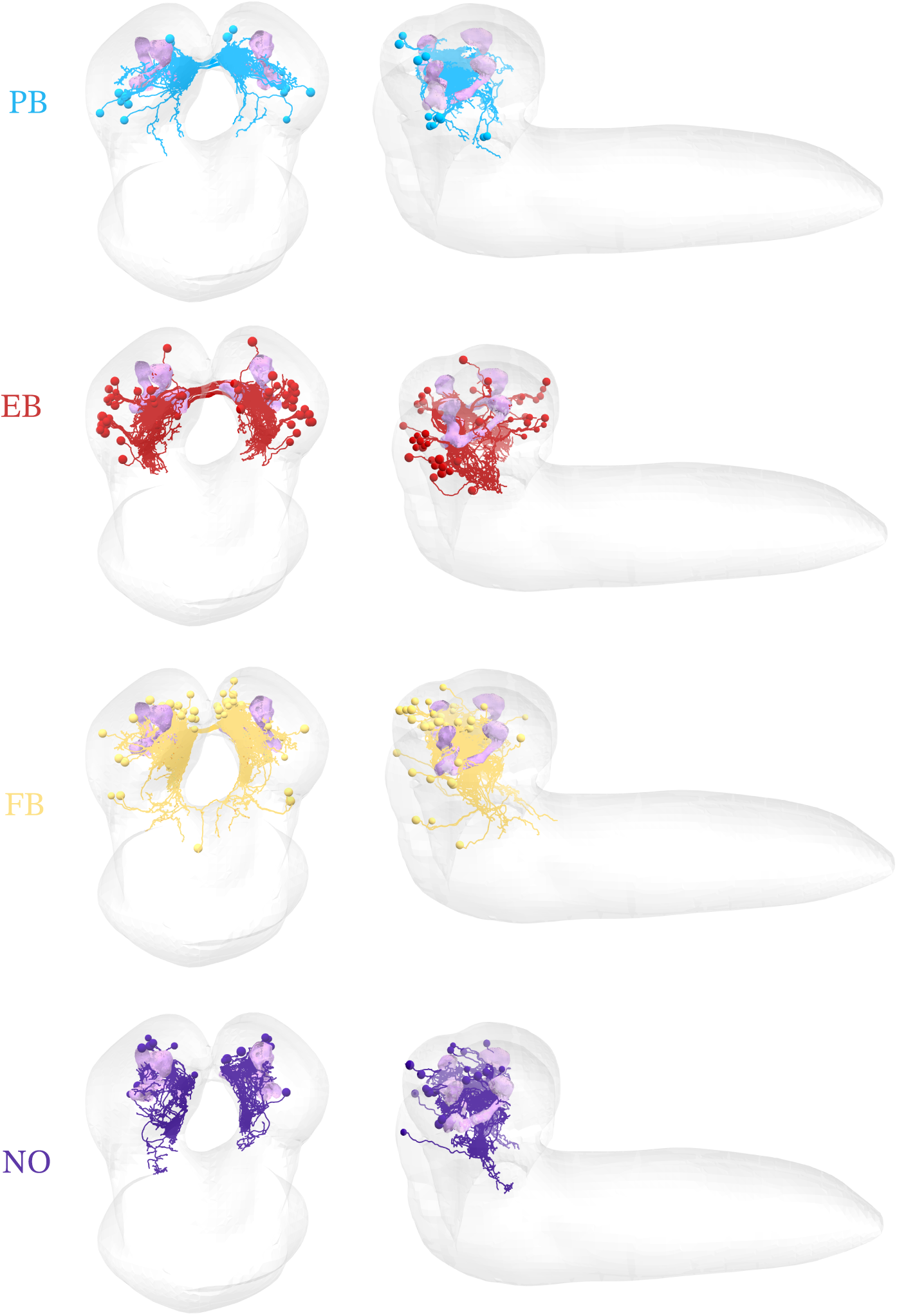
Larval CX neuron boutons define the volumes of the 4 main CX neuropils. Posterior and lateral views of neurons that contribute boutons (“.b”) to each of the 4 neuropils. In concordance with the adult, the PB is the most dorsal and posterior neuropil of the whole bra9in; the FB occupies an intermediate medial position; the EB is the most medial anterior neuropil (same dorso-ventral level as the mushroom body medial lobe but even more anterior); and the NO are medial and ventral. Note that the white medial space in the larval cartoon is truly devoid of brain tissue: the larval CX neuropils line the medial boundary of the brain hemispheres, with the pharynx occupying most of the intermediate space.

In addition to visual input, the adult PB also integrates olfactory inputs [38], suggesting that spatial navigation is not unimodal but integrative across multiple sensory modalities.

In searching for the larval PB, we expected two sets of neurons: columnar and horizontal. In larva, four CX lineages contribute columnar neurons, a subset of which position their dendrites at a posterior-dorsal location. We could not find a CX lineage that would contribute horizontal fibers at a posterior-dorsal location necessary to intersect and synapse onto the dendrites of the PB columnar neurons, but we found a larval lineage (DALv1) whose axons are bilateral and project to the appropriate area, and is spatially adjacent and developmentally related to another central complex lineage (DALv23). This suggests that neurons from non-CX lineages may be recruited temporarily during the larval period into CX neuron types, in a pattern reported so far for the mushroom body (see Discussion; [31]).

##### Larval PB horizontal fibers

Among neurons of the larval DALv1 lineage, 4 left-right pairs (named HF-PB for “Horizontal Fiber PB”) project their axons bilaterally and across the dendrites of the columnar neurons. 3 of the 4 pairs present an unusual axon configuration: first, they project contralaterally to drop their first output synapses, with the axon then backtracking to cross the midline a second time to return back to the same ipsilaterally corresponding location to again drop presynaptic sites (Fig. 3).

**Figure 3:**
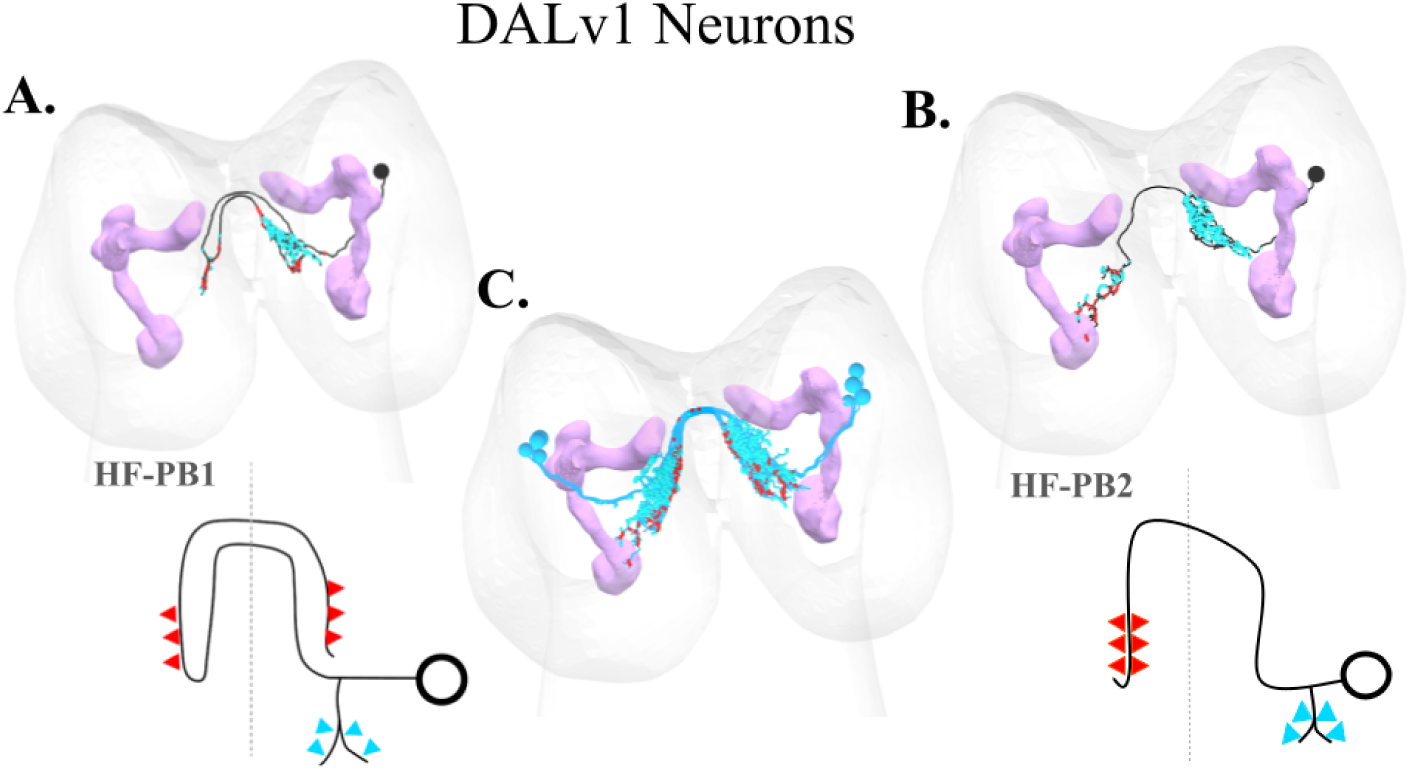
PB Horizontal Fibers (HF-PB). Two types of horizontal fiber neurons are shown, each with a morphological reconstruction (top) and a simplified schematic (bottom). The mushroom body is shown in magenta. Input synapses (dendrites) are indicated by blue arrows, while output synapses (axons) are indicated by red arrows. Panel **A.** shows HF-PB1 neurons, characterized by a double-crossing axon; panel **B.** displays HF-PB2 neurons, with outputs projecting only contralaterally; and panel **C.** the four bilateral pairs of HF-PB neurons shown together.

This peculiar axon configuration is unique among all neurons of the entire brain of the larva ([30]) and suggestive of a delay line for comparing left-right sensory inputs.

The 4th pair of HF-PB (DALv1) neurons first drops presynaptic sites ipsilaterally and then its axon crosses the midline until reaching the corresponding contralateral location to synapse again (Fig. 3).

The presynaptic outputs of all 4 pairs of HF-PB (DALv1) neurons are symmetric, in that they contact the same homologous pairs of left-right neurons which are predominantly neurons of the columnar system from the DM1-DM4 lineages. The axons of these 4 pairs of HF-PB neurons are tiled dorso-ventrally, falling into two bilaterally symmetric groups which we interpret as defining 2 + 2 bilaterally arranged PB compartments, each innervated by 2 pairs of axons (Fig. 3).

The dendrites of these 4 pairs of HF-PB (DALv1) neurons are ipsilateral and dorsal, receiving polysynaptic inputs from vision and olfaction, in a pattern similar to the PB horizontal fibers in the adult ([38]). These multi-sensory inputs to HF-PB are mediated by Convergence Neurons (CN-53 and CN-54, among others (Fig. 3) that integrate inputs from both MBONs (MB Output Neurons) and from the Lateral Horn (LH), such as olfactory, visual, and gustatory PNs among others [55]). Therefore, sensory inputs arriving to the larval PB will have been modulated or gated by previously established associative memories.

Three more larval lineages contribute HF-PB neurons: the BLAvm (neuron SLR), the BLAd-OL (neuron MB2ON-63), and the DPMm2 (neuron ADC1), with one pair of neurons each.

MB2ON-63 presents a bilateral axon similar to the 4th pair of HF-PB (DALv1), and an ipsilateral dendrite that integrates inputs from multiple convergence neurons (CN-54, CN-9), MBONs (MBON-c1, MBON-d1), and sensory PNs (olfactory mPN B1, visual PVL09).

SLR presents a contralateral axon that spans both the FB and PB, and an ipsilateral dendrite that integrates inputs from MB2ON-63, CN-9, and the FB intrinsic neuron MB2ON-175 (see below), in addition to a long tail of weaker input from ascending neurons relaying information from the somatosensory system.

ADC1 presents an ipsilateral dendrite projecting to the posterior brain and a bilateral axon circumscribed to the PB. These neurons integrate inputs from multiple sensory PNs (mPN B2, mPN 5 (also known as mPN BAmd1-g), mPN A1, and weakly from others), a convergent neuron (CN-35), MB2ON-94, and a few others weakly. Their axons target primarily HF-PB neurons (DALv1), but also convergent neurons strongly (CN-56, CN-15, CN-35).

##### Larval PB columnar neurons

In the larva, the columnar system consists of neurons from 4 central complex lineages (DPMpm1, DPMpm2, DPMm1 and CM4) that also generate the columnar neurons of the adult (DM1, DM2, DM3 and DM4, correspondingly: these are the same lineages with different names for the larva and the adult for historical reasons). Larval columnar neurons present small, narrow dendrites circumscribed within the 2 + 2 compartments defined by the axons of the HF-PB (DALv1) neurons, from whom they recieve synapses.

Among the columnar neurons, a subset project their axons directly to the NO (Fig. 4A), and another subset project directly to the larval EB (Fig. 4B).

**Figure 4:**
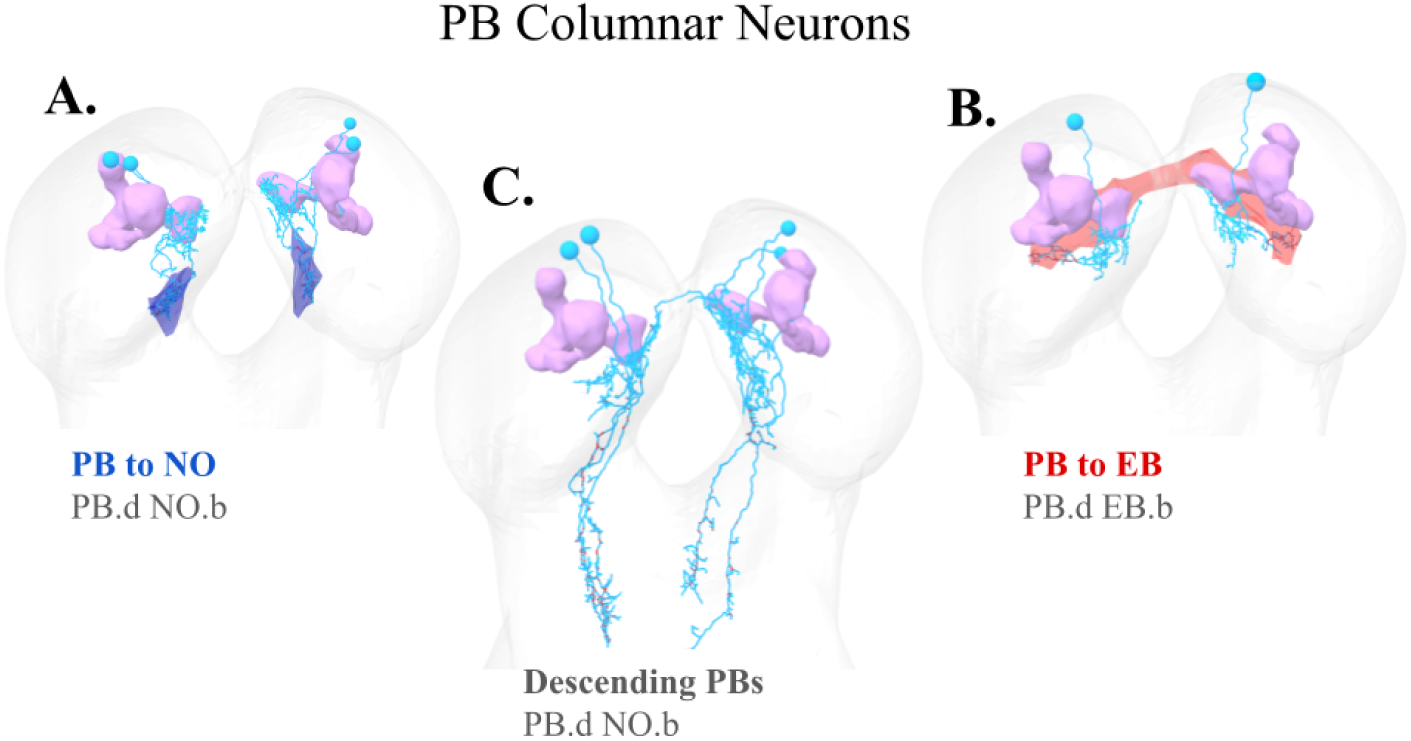
PB columnar neurons. The three types of columnar neurons are shown. The mushroom body is shown in magenta. **A.** Neurons projecting from the PB to the NO (PB.d NO.b), with the NO neuropil shown in dark blue. **B.** Neurons projecting from the PB to the EB (PB.d EB.b), with the EB represented by the red structure beneath the mushroom body. **C.** Descending PB neurons projecting toward the SEZ or the VNC.

We did not find among the larval columnar neurons any whose axon would project to more than one CX neuropil, despite such types being common in the adult [36–38]. Beyond the canonical columnar neurons projecting to other CX neuropils, we found some whose axons descend directly to the SEZ or nerve cord (Fig. 4C).

#### 3.1.2 Ellipsoid Body (EB)

The adult Ellipsoid Body (EB) is a ring-shaped structure situated between the Fan-Shaped Body (FB) and the Mushroom Body horizontal lobes, facing anterodorsally. The EB is made of two main types of neurons: Ring-neurons (RNs; derived mainly from the EBAa1/DALv2 and LALv1/BAmv1/2 lineages) that spread their axons across the length of the EB, and reciprocally connected Wedge neurons (EPGs, derived from the DM4 lineage) that divide the EB into 16 compartments (the wedges) [37]. The EPG (Wedge) neurons are excitatory and synapse with each other, as well as reciprocally with the Ring neurons, which are inhibitory [8, 38].

The adult EB circuit has been modeled as a ring attractor [7] to, in concordance with its anatomy, reproduce *in silico* the observed “bump” of neural activity in the form of a sole active wedge in the *Drosophila* adult EB [56].

EPG (Wedge) neurons form 16 wedges around this ring, and project to both hemispheres of the PB, where they connect to two sets of columnar neurons (PEG and PEN) that project back to the EB, forming recurrent loops. The anatomical offset between EPG and PEN neurons is key to how the fly head direction system translates angular motion into an updated position of the activity bump in the ring attractor [7].

In the adult, the EB receives visual inhibitory GABAergic inputs, via two parallel pathways for distinct visual information: 1. Ring neurons that map the visual environment of the fly; 2. tangential neurons that take in information about body rotations and translational velocity. The latter receive input in the LAL and output to the NO. Mechanosensory input also enters the CX via the second-order projection neurons to the EB. These neurons code head direction; some proprioceptive input has also been observed [38]. It receives strong inputs from PB, NO and the LAL, and outputs onto the PB.

##### Larval EB Wedge neurons

In the larval brain, we found a group of 8 pairs of reciprocally synaptically connected neurons from larval lineage DALcl12, known to produce Wedge neurons in the adult, so we named them Wedge neurons. Both their dendrites and axons are very small, and tiled medio-laterally, defining 8 compartments with one single neuron pair contributing to each (Fig. 5C). We found an additional pair of neurons from larval lineage DALv23 (which produces ring neurons in the adult) with the same connectivity pattern as Wedge neurons (Fig. 5B).

**Figure 5:**
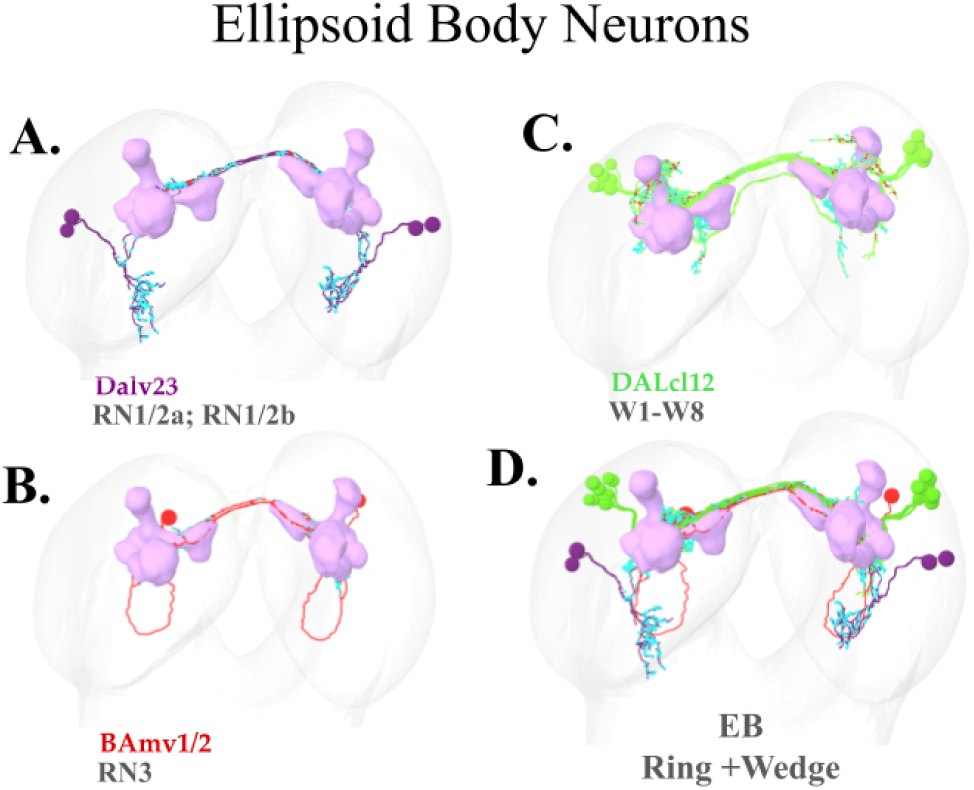
Ellipsoid Body Neurons. Ring and Wedge Neurons with their corresponding lineages. The mushroom body is shown in magenta as a landmark in the larval brain. **A.** Ring neurons of lineage Dalv23, namely RN1/2a and RN1/2b; **B.** Ring neuron RN3 originating from lineage BAmv1/2; and **C.** The 8 EB wedge neurons (W1-8) originating from DALcl12 lineage. Panel **D.** depicts all these neurons stacked together to form the EB

While Wedge neurons make both axo-dendritic and axo-axonic synapses onto each other (like the AL Picky LNs do; [27]), their dendrites integrate inputs from a variety of sources, including MBONs (MBON-e1, MBON-e2, MBON-g1, MBON-g2, MBON-b3, MBON-m1, and weak inputs from more), convergence neurons (CN-14, CN-25, CN-62, CN-38), and MB feedback neurons (FBN-13, FBN-19, FBN-21), indicating that the EB is intimately and intricately involved with the MB (see below). Wedge neurons, like the adult, receive further axo-axonic synapses from Ring neurons (see below; RN1/2, also known as FBN-20; and RN3).

##### Larval EB Ring neurons

We found one pair of neurons (RN3) of the larval BAmv1/2 lineage (corresponding to adult lineage LALv1 known to generate ring neurons) that receive visual input via PB columnar neurons, and reciprocally synapses axo-axonically with larval Wedge-neurons (Fig. 5A). Their axons are fully contained within the space defined by the wedge neurons. Similarly, we found two pairs of neurons (RN1/2) from the larval DALv23 lineage (adult EBa1 lineage, also known to generate ring neurons), synaptically connected like RN1/2 (Fig. 5B). We classified all of these as larval EB Ring neurons.

##### Sensory inputs to the larval EB

In the adult, the EB is known to receive visual inputs via a polysynaptic pathway (see Fig. 6 in [38], and Fig. 4 in [57]). In the larva, we find that the main visual PN for negative phototaxis, PVL09 [58], synapses axo-dendritically onto MB2ON-187, which relays sensory inputs from visual and other sensory modalities axo-axonically onto larval Wedge neurons and Ring neurons (Fig. 7). MB2ON-187 is named so for being postsynaptic to MBONs (MBON-c1 in particular; [55]). Through MB2ON-187, multiple sensory PNs (PVL09, mgPN 7) and many neurons postsynaptic to MBONs (MB2ONs) synapse onto Wedge neurons (see Suppl. Connectivity Matrix). The latter indicate, again, that MBONs modulate or filter sensory inputs converging onto the EB.

**Figure 6:**
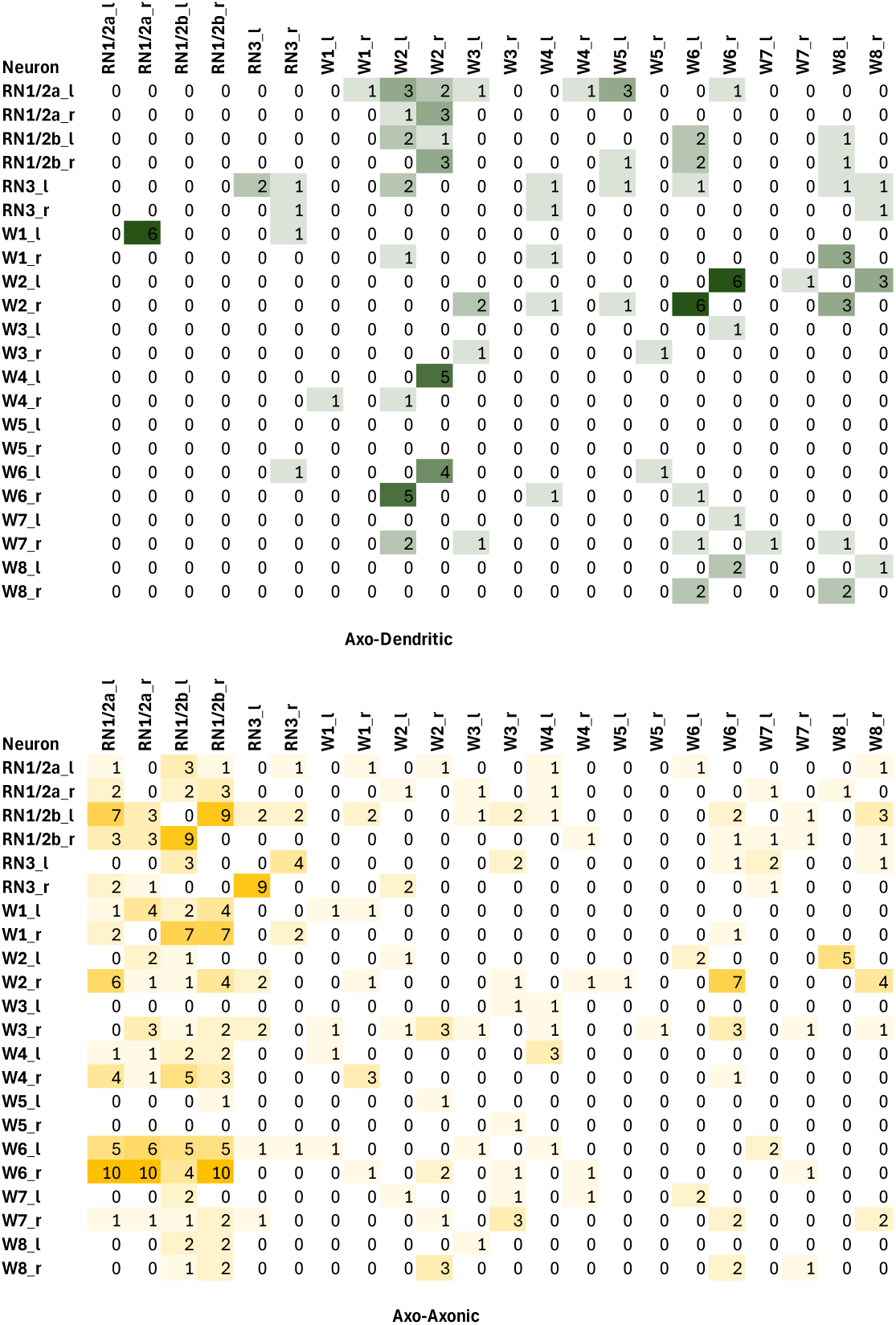
EB Neurons Connectivity. Both tables display the connectivity patterns among ring and wedge neurons in the ellipsoid body (EB). Each neuron is separated into left (_l) and right (_r) counterparts to represent lateralization. **Upper table:** axo-dendritic connections (color-coded in the green spectrum). **Lower table:** axo-axonic connections (color-coded in the yellow spectrum).

**Figure 7:**
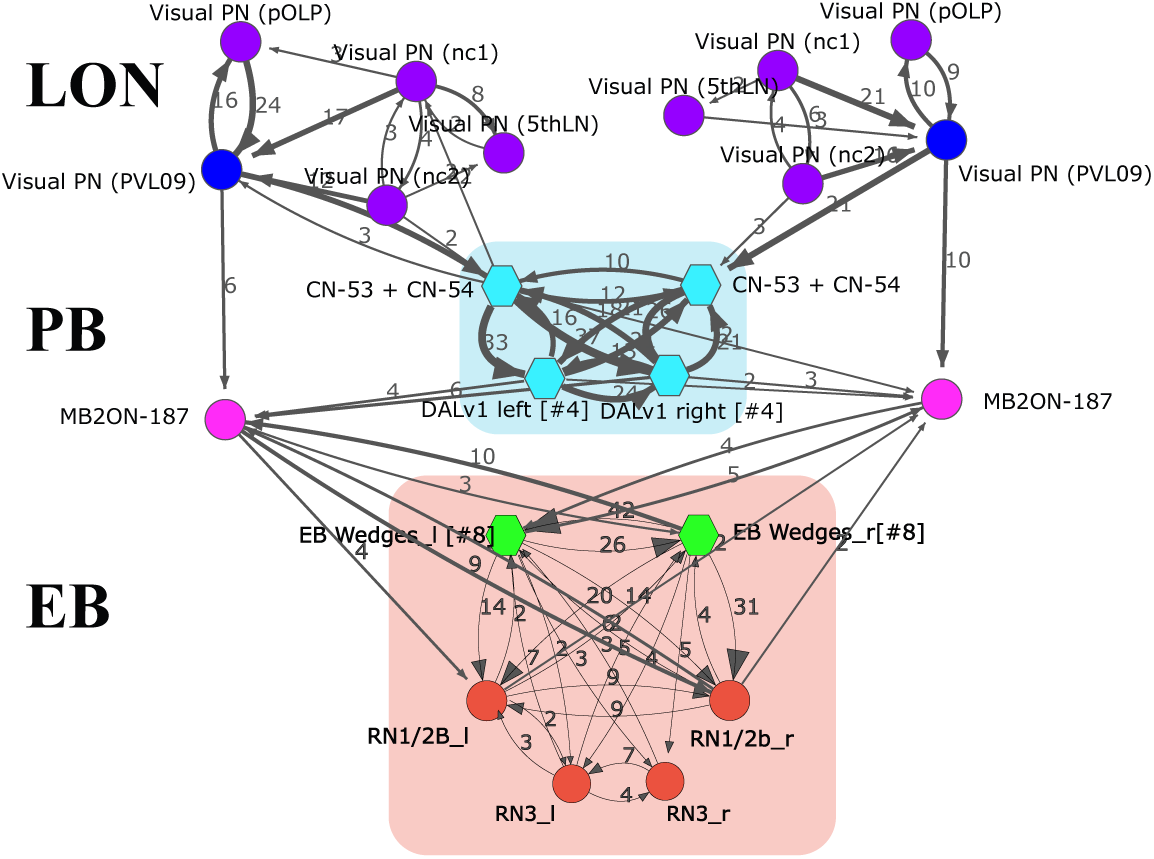
Visual Input into the EB. Input from visual Projection Neurons (Visual PN) to the Ellipsoid Body (EB) via two convergence neurons (CN53, CN54), a pair of PB DALv1s and a second-order Mushroom Body output neuron (MB2ON-187).

#### 3.1.3 Fan-shaped Body (FB)

The adult FB is a bilaterally symmetric neuropil posterior and dorsal to the EB, with well-defined horizontal and vertical components: 6 horizontal layers stacked dorso-ventrally that are defined by distinct sets of horizontal neurons (FB tangential neurons); and 9 vertical columns stacked medio-laterally are defined by column-specific columnar neurons. Both horizontal and vertical neurons innervate the FB in a layer- and column-restricted manner [59]. As the biggest CX neuropils, a large variety of lineages contribute to the FB (see Table 1).

**Table 1:**
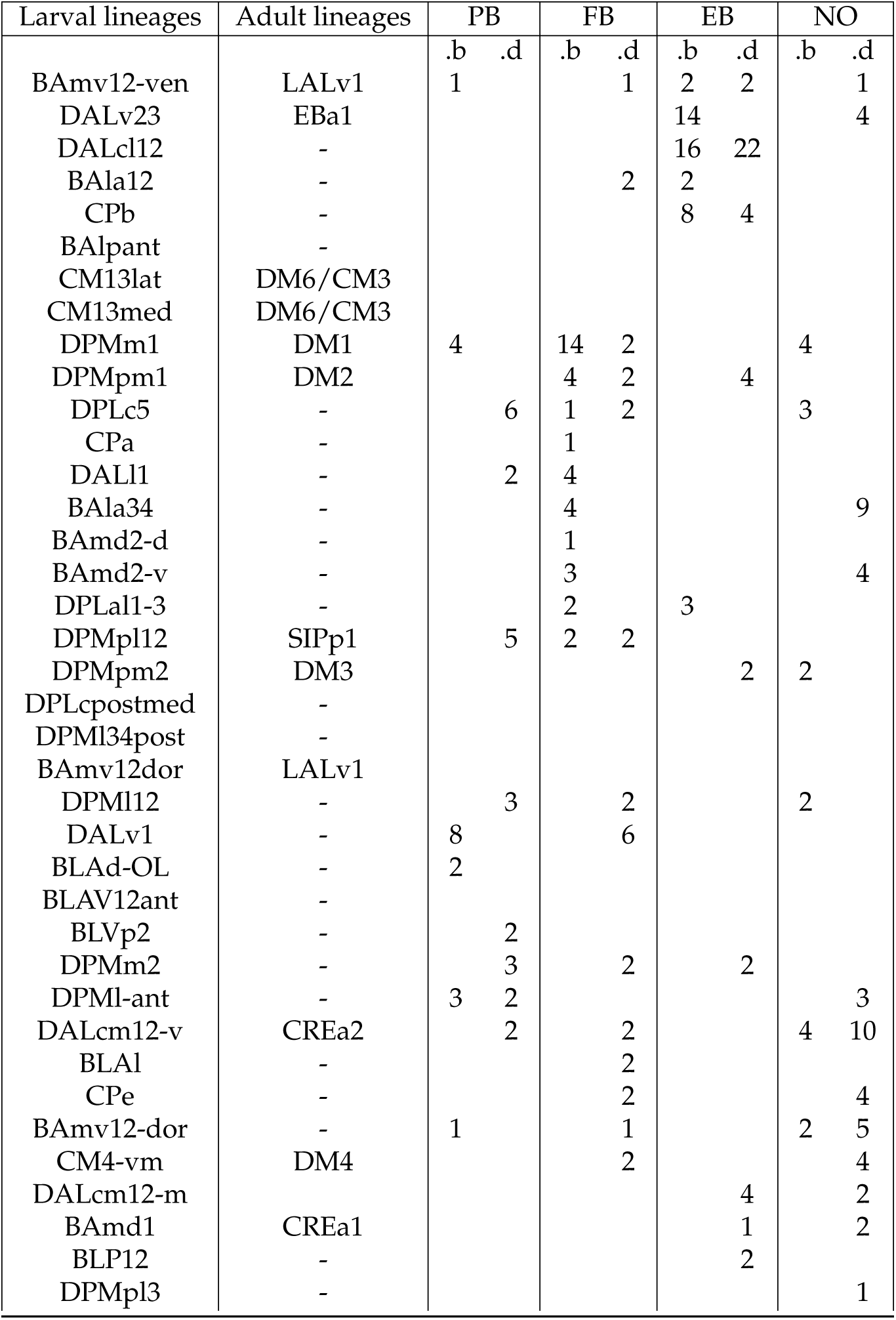
Central complex lineages. Larval CX lineages are listed in the first column. If a lineage also contributes to the adult CX, the corresponding adult nomenclature [53] is shown in the second column. Columns 3–6 indicate the number of neurons from each lineage that contribute axons (“.b” for boutons) or dendrites (“.d”) to individual neuropils: PB, FB, EB and NO.

In the adult FB there are 2 types of FB tangential cells: (1) neurons that relay the presence of an attractive odor to the FB, originating in the MB or the LH (learnt or innate valences; [38]); (2) neurons that relay sleep drive to the FB, whose activity is mandatory for sleep initiation [12].

The adult FB columnar neurons, or columnar input cells, are known as PFN (PB-FB-NO; [37]), with dendrites in the PB and axonic outputs on both the FB and NO. There are 5 main types [38], with their arbors tiling the PB glomeruli, the FB layers, and the NO layers.

The third class of adult FB cells are interneurons with dendrites and axons within the FB. Of these there are 2 main types: FB intrinsic neurons whose arbors lay entirely within the FB, and FB mixed arborisation neurons with additional axonic branches outside the CX and sometimes dendritic branches in the PB [37].

A key feature of the the adult FB is strong innervation by MB Output Neurons (MBONs) [38, 60]. In addition, the axons of dopaminergic neurons driven by visual inputs innervate the FB [61].

In the larva, we found a number of putative FB horizontal/tangential cells originating in lineages known to contribute neurons to the adult FB. Characteristically, most present a bilateral axon closely wrapping around the midline (hence often informally named “horseshoe”), and an ipsilateral dendrite positioned within the superior protocerebrum (dorsal anterior neuropil) where they integrate numerous inputs from MBON axons. Among the many neurons with dendrites within this very medial neuropil, we find neurons from lineages known to contribute to the adult FB (larval lineages DPMpm1, CM13, DPMpl12 and DPMpm2; see Table 1) and whose axons project to the putative larval NO, EB, PB and LAL neuropils.

##### Larval FB tangential cells

We found a diverse and large collection of larval FB tangential cell types. All types contribute synapses rather specifically to the FB and are hence labeled as FB.b, a defining feature across them all. The somas originate in a number of larval lineages whose adult homologs are known to contribute neurons to the adult FB (see above), but also a few from the nerve cord, projecting their ascending axons directly to the FB such as the pCC neurons (a cell type first reported in grasshopper [62] and later in the *Drosophila* embryo [63]). The axons of the pCC project to the ipsilateral FB. The set of ascending neurons isn’t complete, for the ventral nerve cord (VNC) remains incompletely mapped.

Among all neurons projecting their axons to the FB, many are brain neurons that do so with a typical bilateral horseshoe (’hs’) axon, and hence we named then hs-FB, with a numeric label for each subtype.

The DM1-DM4 lineages that contribute to the adult CX also generate a number of neurons in the larva (larval lineages DPMm1, DPMpm1, DPMpm2, and CM4-vm, respectively). Targeting the FB, there are several major subtypes. First, 6 pairs of neurons (hs-FB.1) with a bilaterally symmetric axon that tightly lines the medial surface in a horseshoe shape, and with a smallish dendrite dorsal and slightly lateral to the FB, within the medial side of the superior proto-cerebrum. Second, 2 pairs of neurons (hs-FB.2) also with bilaterally symmetric axons but that target the more posterior-lateral FB and beyond into the larval PB, with also ipsilateral dendrites that extend anteriorly and ventrally along the neuraxis. Third, a pair of neurons (hs-FB.3) with bilateral axons placed towards the most posterior end of the FB and perhaps beyond, into the lateral side, and with small ipsilateral dendrites.

The axons of all 6 pairs of hs-FB.1 neurons define the ventral layer of the FB, and the axons of the hs-FB.2 the dorsal one, which is shorter and more anterior. The hs-FB.3 axons project in between the other two types but also cover the posterior-dorsal area that hs-FB.2 doesn’t (Fig. 8).

**Figure 8:**
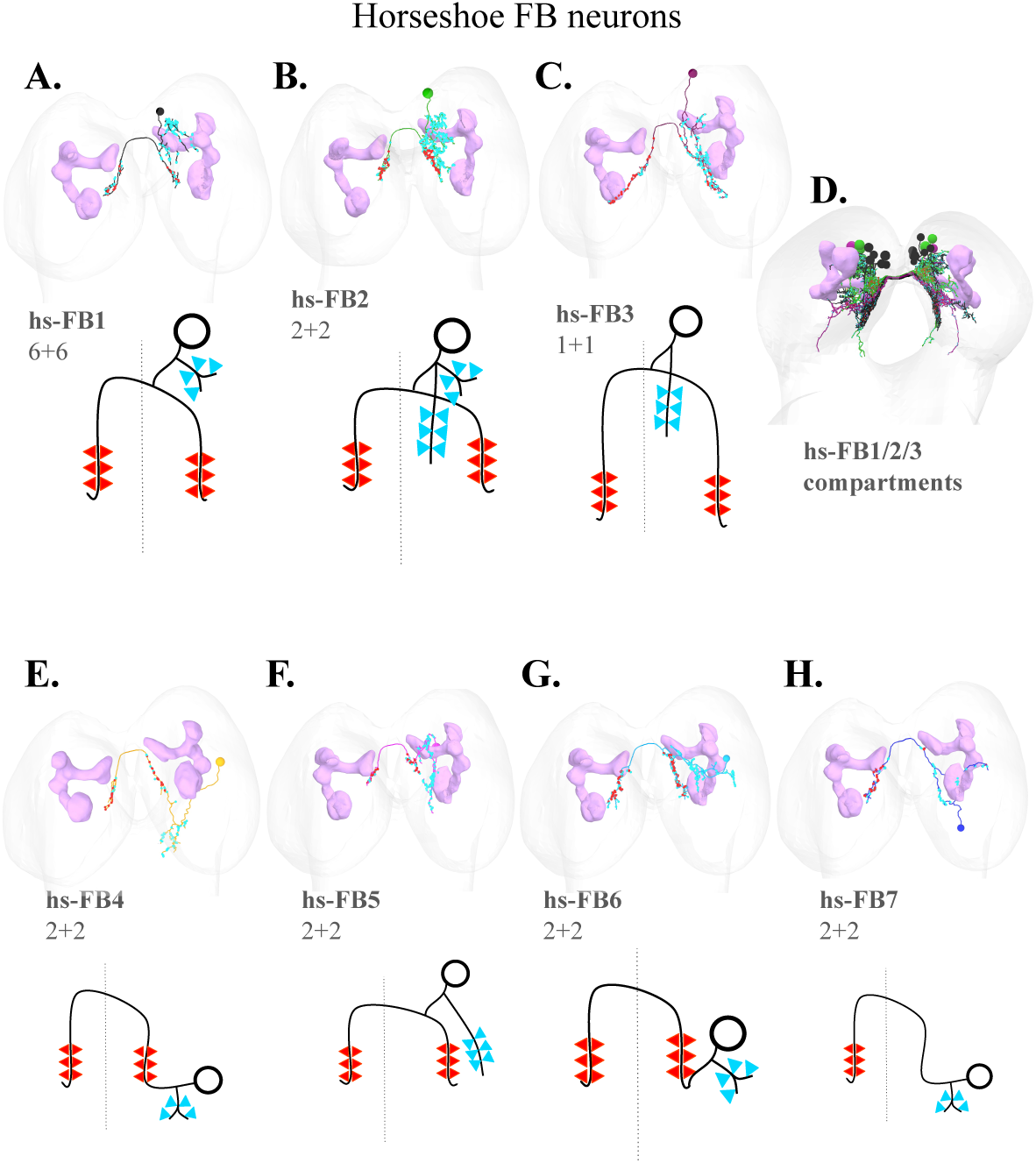
**FB tangential cells,** also known informally as “horseshoe” neurons for the shape of their bilateral axons. The seven hs-FB subtypes are shown, each with a morphological reconstruction (top) and a simplified schematic (bottom). Input synapses (dendrites) are indicated by blue arrows, and output synapses (axons) by red arrows. The mushroom body is shown in magenta. Neuron counts (left+right) are given below each schematic. **A–C.** Individual examples of hs-FB.1, hs-FB.2, and hs-FB.3. **D.** Overlay of hs-FB.1 through .3, illustrating the FB compartments they define. **E–H.** Individual examples of hs-FB.4 through .7.

The larval BAla34 lineage generates sensory PNs, of which two pairs (hs-FB.4) present the typical bilateral axon and their dendrites collect inputs from the ventral posterior brain where ascending PNs from the somatosensory system (mostly mechanosensory) project their axons. One of the pairs is also NO.d (collects inputs from NO.b neurons).

The larval BAmd2 lineage contributes two pairs of neurons (hs-FB.5) with a bilateral axon and dendrites that cover a region medial but lateral to the FB, and also far more ventral.

The larval DALl1 lineage contributes two pairs of neurons (hs-FB.6) with a bilateral axon and dendrites integrating inputs from the intermediate and lateral areas to the FB, primarily from neurons postsynaptic to ascending mechanoreceptive neurons like A00c [40]. A number of neurons presynaptic to hs-FB.6 are also labeled as FFNs and MB2INs, meaning they converge onto MB DANs, MBINs and OANs [64].

The larval CPa lineage contributes a pair of neurons (hs-FB.7) with a bilateral axon and dendrites that cover the most posterior end of the FB, or right outside, plus sparsely a space lateral to the FB, collecting inputs from polysynaptic pathways originating in somatosensory mechanoreceptive neurons.

Finally, the DPLal1-3 lineage contributes a pair of small neurons (ip-FB.1) with an ipsilateral-only axon targeting the most posterior lateral side of the larval FB, and with dendrites more lateral that integrate inputs from a variety of neurons, some of which are NO.d.

##### Larval FB columnar neurons

By definition, all larval FB columnar neurons present dendrites in the FB (are FB.d), and are postsynaptic to FB.b neurons.

In addition to generating FB tangential neurons, the DM1-DM4 lineages also contribute 5 pairs of FB columnar neurons. These neurons (FB.d.1) are characterized by ipsilateral-only dendrites largely circumscribed to the FB (except for the Bushy neuron from the DPMpm1/DM3 lineage), and projecting their ipsilateral-only clutchy axons solely the NO (so they are FB.d-NO.b). Therefore, FB.d.1 are specialized in relaying information from the FB to the NO hemilaterally. Among FB.d.1 we find CN-34, MB2ON-201 and MB2ON-204 [55].

The DM lineage (larval CM4-vm lineage) further contributes 1 pair of neurons (FB.d.4) with ipsilateral dendrites in the posterior FB and beyond into an area adjacent to the LAL medially, and a contralateral axon that descends into the VNC, dropping presynaptic sites from the SEZ to the most posterior abdominal segments. Within the brain, FB.d.4 synapses onto some NO.d neurons (e.g., MB2ON-200) and also onto CN-28 [55].

The MB neuroblasts contribute 3 pairs of early-born, non-Kenyon cell neurons (the ‘ni’ for “non-intrinsic”), named FB.d.2, with ipsilateral dendrites in the FB and a contralateral axon targeting exclusively the LAL.

Then there are a collection of single pairs of neurons, each contributed by a different neuroblast, with dendrites in the FB and their axons targeting widely divergent neuropils, across the brain and also into the SEZ and VNC.

The larval BLAv12 lineage contributes 1 pair of neurons (FB.d.3) with ipsilateral dendrites in the FB and also more medially, and ipsilateral axons targeting exclusively the LAL.

The larval DALcm12-v lineage (adult CREa1) contributes 1 pair of neurons (FB.d.5) with ipsilateral dendrites in the FB and in the area adjacent but medial and ventral to the LAL, projecting their axons ipsilaterally from the NO to the medio-ventral brain and dorsal SEZ. The axons synapse onto a large number of NO.d neurons.

The BAmv12-dor larval lineage contributes 1 pair of neurons (FB.d.6) with compact ipsilateral dendrites within the FB, and a compact short ipsilateral axon immediately dorsal on the PB. The FB.d.6 hence serve as a direct relay from the FB to the PB, synapsing onto hald a dozen PB.d neuron types.

The BAla12 larval lineage contributes 1 pair of neurons (FB.d.7; also known as FFN-6, [55]) with compact ipsilateral dendrites in the FB, and an ipsilateral axon that targets the superior anterior protocerebrum where it synapses profusely onto the dendrites of DANs and OANs of the MB (DAN-i1, DAN-j1, DAN-c1, OAN-g1). Hence, FB.d.7 serves as a relay between the FB and the MB input neurons, conveying teaching signals for associative memories.

The BLAl larval lineage contributes 1 pair of neurons (FB.d.8) with ipsilateral dendrites within the FB but also extensively lateral to the FB, and with peculiarly curving axons that follow the neuraxis, closely apposed to the dendrites of some hs-FB subtypes (hs-FB.1, hs-FB.2, hs-FB.3) and synapsing onto them.

The DPMl2 larval lineage contributes 1 pair of neurons (FB.d.9, also known as FB-LAL1) with ipsilateral dendrites on the posterior hemilateral FB (with inputs from hs-FB.2, hs-FB.5, hs-FB.6) and also reaching into the PB (with inputs only from ADC1), and a contralateral axon targeting the LAL where they synapse onto LAL LNs, EB.d neurons, and multiple descending neurons (DNs) that project to the SEZ and VNC.

Finally, 3 of the 4 pairs of HF-PB neurons (the ones with a doubly crossing axon) are all FB.d, namely, they have dendrites within the FB and receive numerous synapses from FB.d neurons, in particular subtypes hs-FB.2, hs-FB.5 and hs-FB.6 (with further but weaker inputs from other subtypes). This indicates that the PB horizontal fibers not only integrate inputs from multiple sensory systems and gated by MB outputs, but are also modulated by the FB.

##### Larval FB intrinsic neurons

We found two pairs of neurons (MB2ON-125 and eDAN-2) that are both FB.d and FB.b, corresponding to the third type of adult FB neurons: the intrinsic.

MB2ON-125 integrate inputs across many neuron types, from PB.d columnar neurons to LH output neurons. As the MB2ON-125 name indicates (see [55]), it receives direct synapses from MBONs (just MBON-c1, but a strong connection). Fairly large, its dendrites span approximately the entire spatial domain of the putative larval FB. Postsynaptically we find the CSD neuron [27] and ascending neurons from the ventral nerve cord (VNC), among many others such as FFN-31 (so-called feedforward neurons, from the MB perspective, that bring sensory inputs to MB DANs; [55]).

eDAN-2 is a non-MB dopaminergic neuron (DAN) entirely circumscribed to the posterior-lateral end of the FB. See section on DANs of the CX below for details.

#### 3.1.4 Noduli (NO)

The noduli are small, bilaterally symmetric spherical neuropils located medially and ventrally to the FB. In the adult *Drosophila* brain, each hemilateral neuropil is divided in 3 subunits: nodulus 1, 2 and 3 (NO1, NO2, NO3), with NO1 having the highest synaptic density of the three. There are notable variations across insect species, with the number of noduli ranging from two to four per brain hemisphere. While the stacked noduli subunits have been referred to as horizontal layers, no vertical subdivisions have been reported for these structures. Therefore no columnar organization is known.

The NO neurons present a unique morphology featuring compact, clutchy axons, which set them apart from other CX neurons [36, 38] and greatly ease their identification even in the absence of the typical conspicuous anatomical neuropil region present in adult insects. In the adult fruit fly, these neurons primarily originate in the DM1, DM2 and DM3 lineages [35].

In the adult *Drosophila* brain, the NO is interconnected with the EB and the FB, with the latter relaying information to the NO from tangential input neurons via several PB columnar cells such as PEN-neurons (PB-EB-NO; from the head direction system) and PFN-neurons (PB-FB-NO) [37, 38]. The primary NO inputs outside of the CX are from the LAL; these are known as LNO neurons and are suggested to be inhibitory [36, 38]. LNOs send inputs to and receive feedback from columnar neurons. FB tangential neurons make weak reciprocal connections to LNOs and columnar neurons in the NO. All columnar neurons (PFNs and PENs) that synapse onto NO (are NO.b) are recurrently connected to the same LNO (LAL-NO) neurons they receive input from.

In the larva we found a set of 21 neurons (10 pairs plus one unpaired) with highly compact, clutchy axons situated in the posterior ventral area of the brain, from lineages DPMm1 (DM1 in the adult), DALcm12, DPMl12, DPLc5, and BAMv12, as well as a few other larval lineages. We define the volume encapsulating all their clutchy axons as the putative Noduli of the larval *Drosophila*, and labeled all these neurons as NO.b.

Among the 21 NO.b neurons in the larva we find FB.d neurons (four pairs of the FB.d.1 subtype including CN-34), convergence neurons (CN-6, CN-16), neurons postsynaptic to MBONs (MB2ON-37, MB2ON-241), NO-looper (lineage BAmv12dor), MBON-p1 (a multi-compartment MBON of the MB vertical lobe; [28]), and an unpaired neuron (DLPc lineage). Most NO.b neurons integrate inputs from the MB, as their names indicate (see [55]).

Not a single NO.d neuron projects its axon onto any of the other core larval central complex neuropil, but some project to the LAL (Table 2), supporting a role for the NO as an output neuropil of the central complex.

**Table 2:** Monosynaptic connectivity between CX neuropils. Asterisks indicate direct synaptic connections, with row neuropils synapting onto column neuropils. The table summarizes the non-empty intersection sets for annotations in CATMAID such as FB.d with NO.b (i.e., neurons with dendrites in the fan-shaped body and axons in the noduli).

| Neuropil | PB | FB | EB | NO | LAL |
| --- | --- | --- | --- | --- | --- |
| PB |  | * | * | * | * |
| FB | * |  |  | * | * |
| EB |  |  |  |  | * |
| NO |  | * |  |  | * |
| LAL |  |  | * |  |  |
| MBONs |  | * | * |  |  |

**Table 3:**
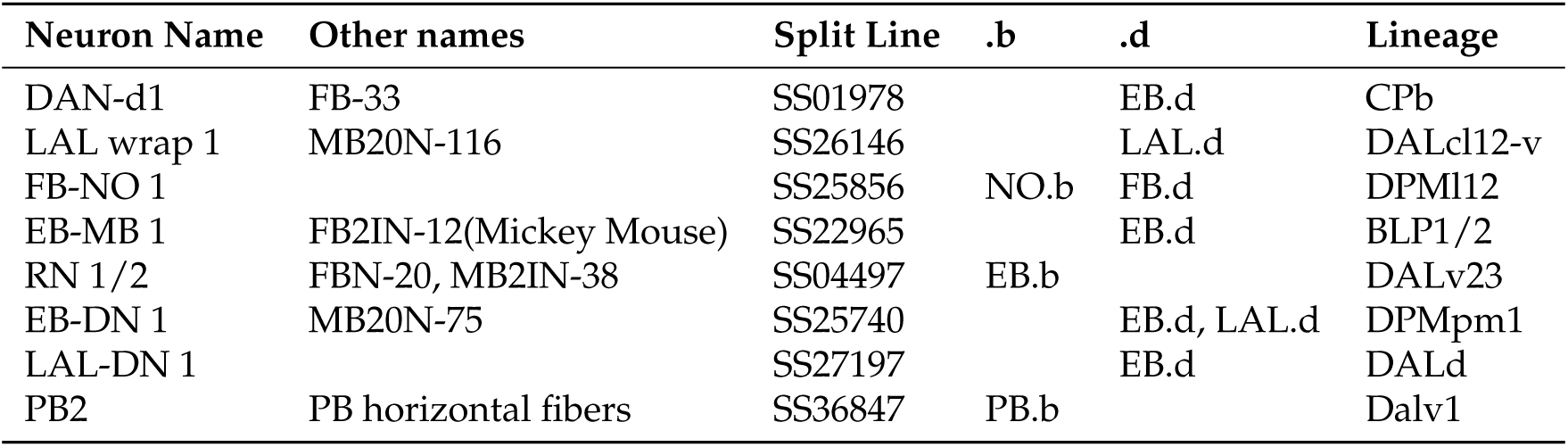
Neurons tested for LOF. Neuron names, corresponding split GAL4 line used for LOF experiments, and lineage information.

### 3.2 Neuromodulation and the CX

In the adult CX numerous neurons express neuromodulators and neuropeptides, and their receptors, particularly in the FB [65, 66]. These modulatory molecules have been associated with the overall level of motor activity and the regulation of sleep [24], among other roles [1]. Here, we list the subset of neurons with assigned neuromodulators that are monosynaptically connected to CX neurons in the larval brain.

The sVUM2mx and sVUM2md are a pair of octopaminergic ventral unpaired medial (VUM) neurons with somas in the SEZ and bilaterally symmetric arbors, which innervate the larval optic neuropil (LON; [29]). These octopaminergic neurons further project their axons into the LAL and also deliver axonic boutons to the EB.

Another octopaminergic VUM (a ladder neuron), named MB2IN-191 [64], innervates the FB bilaterally but not exclusively, with its bilateral axon extending further into nearby medial and posterior areas of the brain.

Given the known role of octopamine in controlling sleep/wake in larvae [10] and the role of the adult CX in modulating motor activity levels and sleep [1, 12], these octopaminergic neurons should be tested experimentally for their potential in mediating a signal for sleep in larvae.

Furthermore, the pair of SP2-1 serotonergic neurons described for the optic lobes [29] present ipsilateral dendrites that integrate inputs from multiple PB.b neurons and also FB.b (hs-FB.6), and then project their axons contralaterally, extending across the PB and into the optic lobe, dropping presynaptic connections onto PB.d neurons and then further posterior and lateral until reaching the LON. These neurons are ideally suited for relaying feedback signals onto the optic lobe to modulate the processing of incoming visual inputs as a function of PB and FB activity.

Another serotonergic neuron, CSD [27], that integrates inputs from olfactory sensory neurons (ORNs) and the superior lateral protocerebrum (an extended area around the LH), and projects back to the antennal lobe (AL) to modulate olfactory LN function [67], integrates strong inputs from the FB intrinsic neuron MB2ON-125.

Taken together, via SP2-1 and CSD neurons the output of the FB may modulate with serotonin both the LON and the AL, presumably to provide context to first-order circuits for sensory processing (vision and olfaction), as has been reported for hunger versus satiation [67].

### 3.3 Dopaminergic neurons (DANs) of the larval central complex

Dopaminergic release outside the MB in the adult fly is known to trigger sleep via dorsal FB (dFB) neurons [68], among other potential roles, depending upon the dopamine receptors expressed in the postsynaptic neurons and the circuits the neurons are part of.

The larval MB houses 7 pairs of DANs [28], with several more DANs scattered across the brain [47]. By flp-out of the TH-GAL4 expression pattern that encompasses all DANs [47] followed by immunohistochemistry, we isolated individual neurons from the TH-GAL4 expression patteern and compared their morphology with EM-reconstructed brain neurons [30]. We identified with reasonable certainty the following 5 eDAN types (eDAN for *external* DAN, i.e., not a MB DAN), comprising 6 neuron pairs.

**eDAN-1**: from the larval lineage DPMpm2, this neuron presents an ipsilateral dendrite in the posterior-dorsal brain, and a bilateral axon exactly overlapping with HF-PB dendrites. eDAN-1 integrates inputs from PB.d neurons like CN-54 (which mediates multi-sensory input integration and relays them to HF-PB, modulated by MB output; see above) but also from multiple sensory PNs such as for temperature (Suckerfish PN, Thermo PN 3 and Thermo PN 5), olfaction (mPN A3), and for unknown sensory modalities (Suckerfish PN 2, from unknown sensory MN-Sens-B2-ACp-21 and -22; [42]), and also from PB.b neurons like MB2ON-63. The axon of eDAN-1 synapses onto multiple PB.d neurons (MB2ON-187, CN-15, CN-54, ADC1, FB-LAL1, and SP2-1) and weakly onto HF-PB neurons.

**eDAN-2**: from the larval lineage CM4-dm, this DAN presents a small, compact ipsilateral dendrite and a contralateral axon on the corresponding contralateral hemisphere right at the posterior end of the FB. eDAN-2 integrates inputs from FB neurons (strongly from FB.d.6, hs-FB.3, hs-FB.1, and weakly from many more). Its axon synapses onto MB2IN-195 (which is presynaptic to EB Wedge neurons), multiple FB neurons (hs-FB.3, hs-FB.1, FB.d.7 (FFN-6)), and weakly onto MB2IN-191 (the octopaminergic VUM of the CX).

**eDAN-3**: also known as MB2IN-139 [55], this neuron is from the larval lineage DPMpl12. Its dendrite is ipsilateral, sitting lateral to the FB but also extending into the LAL. The eDAN-3 axon targets a region dorsal to the LAL, housing the dendrites of many EB.d neurons onto which it synapses, such as MB2IN-114, EB-DN1, MB2ON-17, EB wedge neurons (only W1), and many more neurons such as convergence neurons (CN-30, CN-37, CN-38, CN-41, CN-42, CN-43), MB2ON-256, and a peculiar FB.d.1 neuron (Bushy).

**eDAN-4**: this type consists of two nearly identical neurons that share most inputs but differ only slightly in their outputs, since their axons are juxtaposed but tiled medio-laterally. From the larval lineage DPMpl12, they present an ipsilateral dendrite in the posterior ventral brain and a contralateral axon in the posterior intermediate brain, beyond the limits of the FB proper. Both integrate inputs from numerous EB.d neurons like MB2IN-124, but only eDAN-4l (lateral axon) receives synapse from hs-FB.3. The axons of both eDAN-4 synapse onto some NO.d neurons like MB2ON-248 (whose dendrite has a domain in the NO and another outside where it meets the axon of both eDAN-4) and of other unidentified neurons (lasso-top 3, DALcm12-v descending), but only eDAN-4l synapses onto convergence neurons (CN-9, CN-26), and only eDAN-4m synapses onto hs-FB.5 and hs-FB.7.

**eDAN-5**: from the DPMpm1 larval lineage, this neuron’s dendrite lays posterior to the LAL and projects its axon inside the lateral LAL. The dendrite integrates inputs from EB.d neurons, MB2IN-195, MB2ON-67, and sVUM2mx. The axon targets weakly but axo-dendritically an EB Ring neuron (RN1/2), and more strongly other neurons (MB2ON-67, MB2ON-78, MB2ON-68).

According to [47] there are further non-MB DANs yet to be identified in the larval brain connectome. Note an effort was made to relate the eDANs we could identify to non-MB DAN names in [47] but the resolution mismatch was too great to bridge, except for eDAN-1 which is most likely named “DM3” in [47].

### 3.4 Mushroom Body to Central Complex

In the adult **Drosophila**, the Mushroom Body is known to output onto the Central Complex neuropils through the MB Output Neurons (MBONs). At this stage, adult MBONs primarily target the Fan-shaped Body - tangential neurons from the middle layers (4-6) - and the Noduli - via a direct glutamatergic connection from MBON-30 to LCNOp(LAL–NO) neurons which then target the PFN (PB-FB-NO) neurons [38].

Here, we examine the synaptic relationships between mushroom body input (DANs, MBINs, OANs) and output (MBONs) neurons and the larval central complex neuropils.

#### 3.4.1 A loop between FB and the ‘c’ compartment of the MB

We found that MBON-c1 synapses onto one FB.b neuron, MB2ON-125. Interestingly, DAN-c1 integrates inputs from an FB.d neuron, FFN-6, which in turn integrates inputs from a few neurons that are directly postsynaptic to MB2ON-125, defining a loop between the MB ‘c’ compartment and the FB. No other MBON supplies direct or two-hop inputs onto the FB.

#### 3.4.2 Reciprocal connectivity between EB ring neurons and the ‘g’ compartment of the MB

Four MBONs (a2, b3, g1 and g2) are EB.b (i.e., deliver synapses to the EB; Fig. 9). Interestingly, some of these MBONs are monosynaptically interrelated, namely MBON-g1 and -g2 (which promote approach) are GABAergic neurons that synapse onto MBON-a2 (which promote avoidance) [55]. Furthermore, MBON-g1 and -g2 synapse axo-axonically onto an EB ring neuron, RN1/2 (also known as FBN-20; [55]), a core GABAergic neuron of the EB, which in addition to synapting axo-axonically recyprocally onto EB wedge neurons also synapses onto an EB.d neuron, DAN-g1. Therefore, the output of the MB ‘g’ compartment (MBON-g1 and -g2) modulate, with GABA, the axon of an EB ring neuron (RN1/2), which in turn synapses onto the dendrites of the teaching neuron of the MB ‘g’ compartment (DAN-g1). This circuit configuration defines a close relationship between an associative learning compartment (MB ‘g’) and the EB.

**Figure 9:**
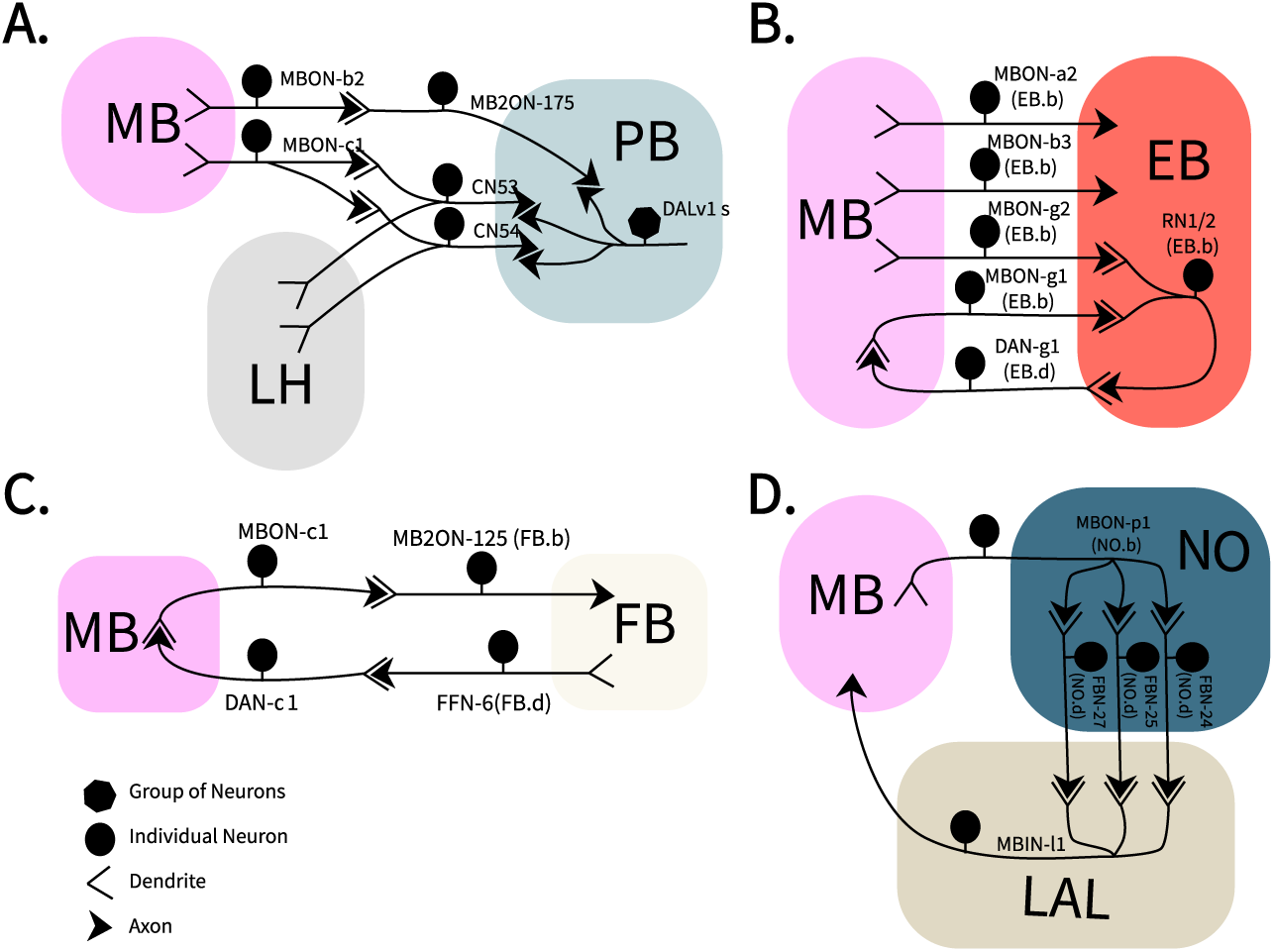
MBONs and DANs inputs into the Central Complex.

Assuming the larval ring neurons RN1/2 are also GABAergic, as they are in the adult [4], the MB ‘g’ compartment is providing disinhibitory input onto the EB, by means of MBON-g1/g2 inhibiting its ring neurons. This disinhibition could be understood as a learning-based gating mechanism over EB wedge neurons, with features of input space expressed in the Kenyon cell population code, by coincident activity with DAN-g1, modulating the activity of the EB wedge neurons that the ring neurons also synapse onto. Taken together, this implies that, if the larval EB is functionally similar to that of the adult EB, there is a component of learning built-in into the internal representation of the direction of movement.

#### 3.4.3 A loop between the NO and the MB vertical lobe via the LAL

MBON-p1 is a multi-compartment MBON of the MB vertical lobe that is an NO.b, (i.e., it delivers output synapses to the NO). At the NO, MBON-p1 synapses onto several NO.d neurons, FBN-24, FBN-25, and FBN-27, which are MB feedback neurons [64] that all project to the LAL and synapse strongly onto MBIN-l1, a MB input neuron targeting multiple compartments of the vertical lobe. This circuit configuration defines a strong tight loop between the MB vertical lobe and the NO via the LAL.

Note that the dendrites of MBIN-l1 are entirely contained within the LAL compartment. In addition, these three feedback neurons also synapse weakly onto DAN-d1.

#### 3.4.4 The MB modulates inputs to the PB

Multiple sensory modalities converge onto the horizontal fibers of the PB (DALv1 neurons) via MB convergence neurons (CN-53, CN-54) and a neuron postsynaptic to MBONs (MB2ON-175). These CNs were previously described as integrating the output of both the lateral horn (LH), which conveys innate pathways, and the mushroom body (MB), for associative memory [55]. Synapses from CN-53, CN-54 and MB2ON-175 onto PB horizontal fiber neurons (DALv1) follow an intriguing pattern of synapting onto either the dendrite or the axon, but largely not both, with specific target choices among the 4 DALv1 neurons. In considering that 3 of the 4 DALv1 neurons (PB horizontal fibers) present an unusual bilateral axon that first deploys output synapses contralaterally and then ipsilaterally (Fig. 3A), the observed pattern of selective axo-dendritic and axo-axonic connections has implications for the modulation of the output of the unusual axons of DALv1 neurons.

In summary, from the perspective of the PB, we find the following circuit architecture: the multi-sensory convergence onto the PB horizontal fibers is directly modulated by the MB, precisely because the neurons that mediate the multi-sensory convergence are themselves CNs (i.e., integrate also MBON synapses in addition to LH inputs): the CN-53 and CN-54. Note CN-54 is in addition a PB.d neuron, integrating inputs from PB horizontal fibers (DALv1 neurons).

Upstream, among various MBONs, MBON-c1 is the most strongly connected to both CN-53 and CN-54, which also integrate inputs from MBON-b1 and MBON-b2. All three are MBONs of the MB peduncle; MBON-c1 has no known function, while MBON-b1 and MBON-b2 promote approach [55].

Of note, the axon of MBON-c1 receives direct presumably inhibitory (GABAergic) inputs from MBON-g1, MBON-g2, and MBON-d1, with all three MBONs participating of circuit loops with other central complex neuropils.

Additionally, neurons directly postsynaptic to MBONs, such as MB2ON-175, in turn directly synapse onto the horizontal fibers of the PB, either axo-axonically or axo-dendritically, in a pattern selective of specific DALv1 neurons.

In summary, not only are navigational decisions such as whether to turn to stimuli or not proceeding not independently per sensory modality but integrated across modalities, as observed before in behavioral experiments [21], but also such integration is modulated by prior memories.

### 3.5 Loss-of-function analysis

We identified genetic driver lines for eight neurons of the larval central complex: EB-MB1, PB2, LAL-DN1, LAL wrap1, FB-NO1, EB-DN1, RN1/2, plus DAN-d1, which is an EB.d (i.e., integrates inputs from the EB). We quantified larval turning behavior following targeted inhibition of each of these driver lines via Tetanus neurotoxin (TNT). Control animals expressed the inactive variant, Impotent Tetanus (ImpTNT). Given that the central complex is classically implicated in visually guided navigation, we used green light stimulation as the sensory stimuli.

We found that expressing TNT did not show significant changes to turning rates for most neurons when compared to controls in response to green light stimulation, with the exception of DAN-d1 (Fig. 10).

**Figure 10:**
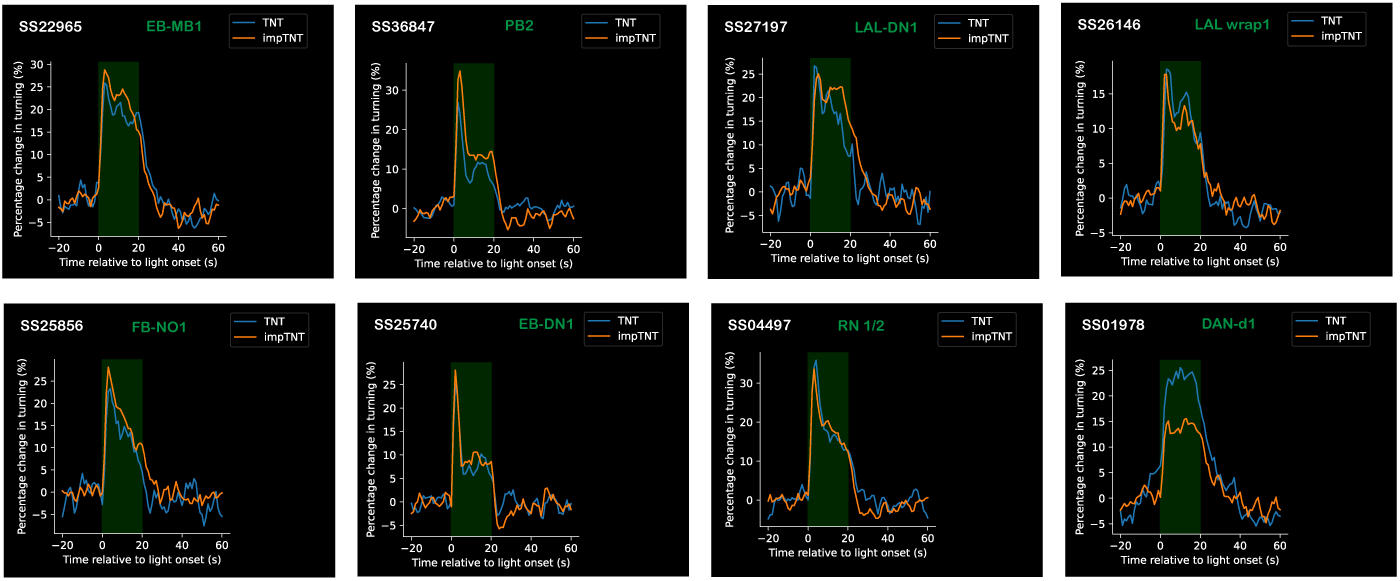
Loss-of-function Behavioral analysis of CX neurons expressing TNT and impTNT(disabled TNT) The name of each split GAL4 line is written in white and their corresponding neuron name in green. Stimulation via green light is highlighted in green from 0-20s. The plots show a time-series of larvae’s turning rates, colored in blue for larvae expressing TNT and orange for larve expressing impTNT

## 4 Discussion

Our inquire into the nature of a fraction of neurons in the remaining *terra incognita* of the larval brain of the fruit fly yielded a series of neurons, neuropils and circuits with undeniable developmental, connectivity, and anatomical similarities to the set of core neuropils of the central complex of the adult fly brain. The key anatomical difference of split brain hemispheres and the very early arrest of neuroblast proliferation in the larvae of Diptera impose strong constraints as to the extent of the anatomical and cellular similarities between the larval and the adult brain.

Notwithstanding, most larval brain neurons survive metamorphosis, help articulate homologous brain neuropils in the adult [69], and participate of neural circuits in the adult brain. For example, the command neurons for backward locomotion [70], and MB MBINs and MBONs [31].

Intriguingly, while we find that larval brain neuronal lineages contribute the same neuronal cell types to the same CX neuropils as their corresponding adult brain instances do, we also find a number of neurons contributing to the larval CX that originate in non-CX lineages. This pattern has been reported before for the MB [31], with some larval MB neurons later on either undergoing apoptosis or, astonishingly, contributing to CX neuropils after metamorphosis. With the MB being an undeniably homologous structure to that of the adult and the larval Kenyon cells making up the bulk of the adult fly brain’s gamma Kenyon cells, the recruitment of non-CX lineages into contributing CX neurons in the larva is perhaps part of a common theme in an animal with a limited neuron budget. The broader context being that the larva of the holometabola is derived [32, 71], an evolutionary novelty that enabled this group of animals to exploit novel ecological niches that, to boot, avoids competition with the adults. In such predicament, it is perhaps not surprising that other larval neuropils similarly to the MB temporarily adopt neurons as their own that will eventually either perish by apoptosis or return to an ancestral cell fate away from the CX.

### 4.1 Navigation, place learning, and memory

The central complex has been associated with place learning (for review see [1]), with the EB in particular being necessary for visual place learning in flies [5]. With fly larvae being capable of local search behavior [72], the structure of the larval central complex and its close interactions with the center for associative memory, the mushroom body, must be examined.

We reported that the ‘c’ compartment of the larval MB is intimately associated with the FB. Intriguingly, MBON-c1 lacks a known role in associative memory to date, with all tests having been performed for olfactory associative memory [64]. Likewise, we discovered a tight loop between the NO and the MB vertical lobe, by means of neurons that also lack a known function in learning: the multi-compartment MBON-p1, whose dendrites span across the MB vertical lobe, and MBIN-l1, a MB input neuron of unknown neurotransmitter signature and function. Both of these MB compartments ought to be examined experimentally for potential roles in navigation or place learning.

With the NO having a role in the adult fly brain controlling the time course of walking activity [73] and in handedness in locomotion [74], and the larval MB vertical lobe mediating aversive memories [64], our report of a tight loop between the output of the MB vertical lobe and modulatory input onto it via the NO may provide the necessary corollary discharge to adjust MB operations to imminent locomotor activity, potentially laterally biased since both MBON-p1 and MBIN-l1 are both entirely ipsilateral-only neurons.

Furthermore, we found that the ‘g’ compartment of the MB, whose MBON-g1 and MBON-g2 promote avoidance [55] and its DAN-g1 is implicated in aversive learning [64], is intimately and recurrently associated with the Wedge and Ring neurons of the EB. Two further MBONs, MBON-a2 and MBON-b3, likewise synapse onto EB neurons. Among these, the function of MBON-b3 remains unknown. Intriguingly, both MBON-a2 and MBON-b3 are rare among the MBONs in having an ipsilateral-only dendrite but a bilateral axon, when the arborization pattern of MBONs is most often the opposite: bilateral dendrites and an ipsilateral axon. The ability of these MBONs to differentially discern sensory inputs between the left and right sides of the body is suggestive of a role in lateralized behavioral responses, such as in taxis.

In loss-of-function experiments, the suppression of DAN-d1 with TNT removes excitatory drive from MBON-d1, which is GABAergic [55]. From the connectome we interpret that a neuron postsynaptic to MBON-d1, MB2ON-63, which is a PB horizontal fiber distinct from DALv1, will then receive less GABAergic input. And therefore, MB2ON-63 receives unopposed excitatory input from visual PNs (PVL09) and other excitatory visual neurons (ChalOLP), and from MBON-c1, all of which are cholinergic [29, 55]. We speculate that, if in wild-type conditions the inhibition from the MB via MBON-d1 was potentially subtracting an expected intensity of light, to perform a temporal comparison, without it the animal would no longer be able to assert where the light has increased or decreased, leading to the need for continuing to sample by head casting or turning.

### 4.2 DANs in the FB and PB

In the adult CX, dFB is a DAN that releases dopamine at the dorsal FB and is capable of triggering sleep [68]. Intriguingly, we found a DAN (eDAN-2) among the FB intrinsic neurons, and two additional DANs (eDAN-4l and eDAN-4m) that synapse onto a number of FB.d, FB.b and NO.d neurons. Identifying genetic driver lines for these neurons will enable testing experimentally their potential role in sleep regulation in larvae.

The role of eDAN-1, projecting to the larval PB, could be related to overall responsiveness and alertness of the animal, since there is a report that DANs projecting to the adult PB mediate increases in aggressiveness [75], and other DANs also projecting to the adult PB decrease sleep [76].

What any of the identified non-MB DANs do awaits the identification of specific genetic driver lines and appropriate experimental setups to study their function.

### 4.3 Conclusion

In conclusion, our interpretation of some of the until now underexamined larval brain circuits as a numerically reduced version of the adult fly central complex is coherent with the evolution of the larval stage in the Holometabola and with the known cell types and overall synaptic connectivity of the corresponding neuropils in adult fly brain, bringing a whole field of study into a life stage of reduced dimensions and numbers of neurons and synapses. Now, with our identification of tight circuit loops between understudied MB compartments and the larval CX, an opportunity opens to examine the neural circuit basis of spatial navigation and place learning in this experimentally tractable animal.

## Acknowledgements

We thank Sam Harris and Kun Wang for their helo with the behavioural assays, and Pedro J. Gómez Gálvez for assistance with immunohistochemistry. We thank the Wellcome Trust (grant 205038/Z/16/Z) and the MRC LMB for funding.

## References

[1] Keram Pfeiffer and Uwe Homberg. Organization and functional roles of the central complex in the insect brain. Annual review of entomology, 59(1):165–184, 2014.

[2] Daniel B Turner-Evans and Vivek Jayaraman. The insect central complex. Current Biology, 26(11):R453–R457, 2016.

[3] Stanley Heinze. Variations on an ancient theme—the central complex across insects. Current Opinion in Behavioral Sciences, 57:101390, 2024.

[4] Ulrike Hanesch, K-F Fischbach, and Martin Heisenberg. Neuronal architecture of the central complex in drosophila melanogaster. Cell and Tissue Research, 257(2):343–366, 1989.

[5] Tyler A Ofstad, Charles S Zuker, and Michael B Reiser. Visual place learning in drosophila melanogaster. Nature, 474(7350):204–207, 2011.

[6] Johannes D Seelig and Vivek Jayaraman. Feature detection and orientation tuning in the drosophila central complex. Nature, 503(7475):262–266, 2013.

[7] Thomas Stone, Barbara Webb, Andrea Adden, Nicolai Ben Weddig, Anna Honkanen, Rachel Templin, William Wcislo, Luca Scimeca, Eric Warrant, and Stanley Heinze. An anatomically constrained model for path integration in the bee brain. Current Biology, 27(20):3069–3085, 2017.

[8] Romain Franconville, Celia Beron, and Vivek Jayaraman. Building a functional connectome of the drosophila central complex. Elife, 7:e37017, 2018.

[9] Stanley Heinze, Ajay Narendra, and Allen Cheung. Principles of insect path integration. Current Biology, 28(17):R1043–R1058, 2018.

[10] Milan Szuperak, Matthew A Churgin, Austin J Borja, David M Raizen, Christopher Fang-Yen, and Matthew S Kayser. A sleep state in drosophila larvae required for neural stem cell proliferation. Elife, 7:e33220, 2018.

[11] Ioannis Pisokas, Stanley Heinze, and Barbara Webb. The head direction circuit of two insect species. Elife, 9:e53985, 2020.

[12] Orie T Shafer and Alex C Keene. The regulation of *Drosophila* sleep. Current Biology, 31(1):R38– R49, 2021.

[13] Yvette E Fisher. Flexible navigational computations in the drosophila central complex. Current opinion in neurobiology, 73:102514, 2022.

[14] Elane Fishilevich, Ana I Domingos, Kenta Asahina, Félix Naef, Leslie B Vosshall, and Matthieu Louis. Chemotaxis behavior mediated by single larval olfactory neurons in drosophila. Current biology, 15(23):2086–2096, 2005.

[15] Sukant Khurana and Obaid Siddiqi. Olfactory responses of drosophila larvae. Chemical senses, 38(4):315–323, 2013.

[16] Shimaa AM Ebrahim, Hany KM Dweck, Johannes Stökl, John E Hofferberth, Federica Trona, Kerstin Weniger, Jürgen Rybak, Yoichi Seki, Marcus C Stensmyr, Silke Sachse, et al. Drosophila avoids parasitoids by sensing their semiochemicals via a dedicated olfactory circuit. PLoS biology, 13(12):e1002318, 2015.

[17] Alex Davies, Matthieu Louis, and Barbara Webb. A model of drosophila larva chemotaxis. PLoS computational biology, 11(11):e1004606, 2015.

[18] Elena P Sawin-McCormack, Marla B Sokolowski, and Ana Regina Campos. Characterization and genetic analysis of drosophila melanogaster photobehavior during larval development. Journal of neurogenetics, 10(2):119–135, 1995.

[19] Zhefeng Gong. Behavioral dissection of drosophila larval phototaxis. Biochemical and Biophysical Research Communications, 382(2):395–399, 2009.

[20] Alex C Keene and Simon G Sprecher. Seeing the light: photobehavior in fruit fly larvae. Trends in neurosciences, 35(2):104–110, 2012.

[21] Ruben Gepner, Mirna Mihovilovic Skanata, Natalie M Bernat, Margarita Kaplow, and Marc Gershow. Computations underlying drosophila photo-taxis, odor-taxis, and multi-sensory integration. Elife, 4:e06229, 2015.

[22] Amy R Poe, Lucy Zhu, Milan Szuperak, Patrick D McClanahan, Ron C Anafi, Benjamin Scholl, Andreas S Thum, Daniel J Cavanaugh, and Matthew S Kayser. Developmental emergence of sleep rhythms enables long-term memory in drosophila. Science Advances, 9(36):eadh2301, 2023.

[23] Jeffrey M Donlea, Matthew S Thimgan, Yasuko Suzuki, Laura Gottschalk, and Paul J Shaw. Inducing sleep by remote control facilitates memory consolidation in drosophila. Science, 332(6037):1571–1576, 2011.

[24] Jeffrey M Donlea. Roles for sleep in memory: insights from the fly. Current opinion in neurobiology, 54:120–126, 2019.

[25] Haluk Lacin and James W Truman. Lineage mapping identifies molecular and architectural similarities between the larval and adult drosophila central nervous system. Elife, 5:e13399, 2016.

[26] Wayne Pereanu, Abilasha Kumar, Arnim Jennett, Heinrich Reichert, and Volker Hartenstein. Development-based compartmentalization of the drosophila central brain. Journal of Comparative Neurology, 518(15):2996–3023, 2010.

[27] Matthew E Berck, Avinash Khandelwal, Lindsey Claus, Luis Hernandez-Nunez, Guangwei Si, Christopher J Tabone, Feng Li, James W Truman, Rick D Fetter, Matthieu Louis, et al. The wiring diagram of a glomerular olfactory system. Elife, 5, 2016.

[28] Katharina Eichler, Feng Li, Ashok Litwin-Kumar, Youngser Park, Ingrid Andrade, Casey M Schneider-Mizell, Timo Saumweber, Annina Huser, Claire Eschbach, Bertram Gerber, et al. The complete connectome of a learning and memory centre in an insect brain. Nature, 548(7666):175–182, 2017.

[29] Ivan Larderet, Pauline MJ Fritsch, Nanae Gendre, G Larisa Neagu-Maier, Richard D Fetter, Casey M Schneider-Mizell, James W Truman, Marta Zlatic, Albert Cardona, and Simon G Sprecher. Organization of the drosophila larval visual circuit. Elife, 6:e28387, 2017.

[30] Michael Winding, Benjamin D Pedigo, Christopher L Barnes, Heather G Patsolic, Youngser Park, Tom Kazimiers, Akira Fushiki, Ingrid V Andrade, Avinash Khandelwal, Javier Valdes-Aleman, et al. The connectome of an insect brain. Science, 379(6636):eadd9330, 2023.

[31] James W Truman, Jacquelyn Price, Rosa L Miyares, and Tzumin Lee. Metamorphosis of memory circuits in *Drosophila* reveals a strategy for evolving a larval brain. Elife, 12:e80594, 2023.

[32] James W Truman and Lynn M Riddiford. The origins of insect metamorphosis. Nature, 401(6752):447–452, 1999.

[33] Nikolaus Dieter Bernhard Koniszewski, Martin Kollmann, Mahdiyeh Bigham, Max Farnworth, Bicheng He, Marita Büscher, Wolf Hütteroth, Marlene Binzer, Joachim Schachtner, and Gregor Bucher. The insect central complex as model for heterochronic brain development—background, concepts, and tools. Development genes and evolution, 226(3):209–219, 2016.

[34] Max S Farnworth, Kolja N Eckermann, and Gregor Bucher. Sequence heterochrony led to a gain of functionality in an immature stage of the central complex: A fly–beetle insight. PLoS biology, 18(10):e3000881, 2020.

[35] Ingrid V Andrade, Nadia Riebli, Bao-Chau M Nguyen, Jaison J Omoto, Albert Cardona, and Volker Hartenstein. Developmentally arrested precursors of pontine neurons establish an embryonic blueprint of the *Drosophila* central complex. Current Biology, 29(3):412–425, 2019.

[36] Tanya Wolff and Gerald M Rubin. Neuroarchitecture of the drosophila central complex: A catalog of nodulus and asymmetrical body neurons and a revision of the protocerebral bridge catalog. Journal of Comparative Neurology, 526(16):2585–2611, 2018.

[37] Tanya Wolff, Nirmala A Iyer, and Gerald M Rubin. Neuroarchitecture and neuroanatomy of the drosophila central complex: A gal4-based dissection of protocerebral bridge neurons and circuits. Journal of Comparative Neurology, 523(7):997–1037, 2015.

[38] Brad K Hulse, Hannah Haberkern, Romain Franconville, Daniel B Turner-Evans, Shin-ya Takemura, Tanya Wolff, Marcella Noorman, Marisa Dreher, Chuntao Dan, Ruchi Parekh, et al. A connectome of the drosophila central complex reveals network motifs suitable for flexible navigation and context-dependent action selection. ELife, 10:e66039, 2021.

[39] Joshua D Mast, Consuelo M De Moraes, Hans T Alborn, Luke D Lavis, and David L Stern. Evolved differences in larval social behavior mediated by novel pheromones. Elife, 3:e04205, 2014.

[40] Tomoko Ohyama, Casey M Schneider-Mizell, Richard D Fetter, Javier Valdes Aleman, Romain Franconville, Marta Rivera-Alba, Brett D Mensh, Kristin M Branson, Julie H Simpson, James W Truman, et al. A multilevel multimodal circuit enhances action selection in drosophila. Nature, 520(7549):633–639, 2015.

[41] Philipp Schlegel, Michael J Texada, Anton Miroschnikow, Andreas Schoofs, Sebastian Hückesfeld, Marc Peters, Casey M Schneider-Mizell, Haluk Lacin, Feng Li, Richard D Fetter, et al. Synaptic transmission parallels neuromodulation in a central food-intake circuit. Elife, 5:e16799, 2016.

[42] Anton Miroschnikow, Philipp Schlegel, Andreas Schoofs, Sebastian Hueckesfeld, Feng Li, Casey M Schneider-Mizell, Richard D Fetter, James W Truman, Albert Cardona, and Michael J Pankratz. Convergence of monosynaptic and polysynaptic sensory paths onto common motor outputs in a drosophila feeding connectome. Elife, 7:e40247, 2018.

[43] Sebastian Hückesfeld, Philipp Schlegel, Anton Miroschnikow, Andreas Schoofs, Ingo Zinke, André N Haubrich, Casey M Schneider-Mizell, James W Truman, Richard D Fetter, Albert Cardona, et al. Unveiling the sensory and interneuronal pathways of the neuroendocrine connectome in drosophila. Elife, 10:e65745, 2021.

[44] Luis Hernandez-Nunez, Alicia Chen, Gonzalo Budelli, Matthew E Berck, Vincent Richter, Anna Rist, Andreas S Thum, Albert Cardona, Mason Klein, Paul Garrity, et al. Synchronous and opponent thermosensors use flexible cross-inhibition to orchestrate thermal homeostasis. Science advances, 7(35):eabg6707, 2021.

[45] Hsing-Hsi Li, Jason R Kroll, Sara M Lennox, Omotara Ogundeyi, Jennifer Jeter, Gina Depasquale, and James W Truman. A gal4 driver resource for developmental and behavioral studies on the larval cns of drosophila. Cell reports, 8(3):897–908, 2014.

[46] Geoffrey W Meissner, Allison Vannan, Jennifer Jeter, Kari Close, Gina M DePasquale, Zachary Dorman, Kaitlyn Forster, Jaye Anne Beringer, Theresa Gibney, Joanna H Hausenfluck, et al. A split-gal4 driver line resource for drosophila neuron types. elife, 13:RP98405, 2025.

[47] Mareike Selcho, Dennis Pauls, Kyung-An Han, Reinhard F Stocker, and Andreas S Thum. The role of dopamine in drosophila larval classical olfactory conditioning. PloS one, 4(6):e5897, 2009.

[48] Shana R Spindler and Volker Hartenstein. The drosophila neural lineages: a model system to study brain development and circuitry. Development genes and evolution, 220(1):1–10, 2010.

[49] Wayne Pereanu, Amelia Younossi-Hartenstein, Jennifer Lovick, Shana Spindler, and Volker Hartenstein. Lineage-based analysis of the development of the central complex of the *Drosophila* brain. Journal of Comparative Neurology, 519(4):661–689, 2011.

[50] Aljoscha Nern, Barret D Pfeiffer, and Gerald M Rubin. Optimized tools for multicolor stochastic labeling reveal diverse stereotyped cell arrangements in the fly visual system. Proceedings of the National Academy of Sciences, 112(22):E2967–E2976, 2015.

[51] Tomoko Ohyama, Tihana Jovanic, Gennady Denisov, Tam C Dang, Dominik Hoffmann, Rex A Kerr, and Marta Zlatic. High-throughput analysis of stimulus-evoked behaviors in drosophila larva reveals multiple modality-specific escape strategies. PloS one, 8(8):e71706, 2013.

[52] Nicholas A Swierczek, Andrew C Giles, Catharine H Rankin, and Rex A Kerr. High-throughput behavioral analysis in c. elegans. Nature methods, 8(7):592–598, 2011.

[53] Nils Eckstein, Alexander Shakeel Bates, Andrew Champion, Michelle Du, Yijie Yin, Philipp Schlegel, Alicia Kun-Yang Lu, Thomson Rymer, Samantha Finley-May, Tyler Paterson, et al. Neurotransmitter classification from electron microscopy images at synaptic sites in drosophila melanogaster. Cell, 187(10):2574–2594, 2024.

[54] Stanley Heinze, Sascha Gotthardt, and Uwe Homberg. Transformation of polarized light information in the central complex of the locust. Journal of Neuroscience, 29(38):11783–11793, 2009.

[55] Claire Eschbach, Akira Fushiki, Michael Winding, Bruno Afonso, Ingrid V Andrade, Benjamin T Cocanougher, Katharina Eichler, Ruben Gepner, Guangwei Si, Javier Valdes-Aleman, et al. Circuits for integrating learned and innate valences in the insect brain. Elife, 10:e62567, 2021.

[56] Johannes D Seelig and Vivek Jayaraman. Neural dynamics for landmark orientation and angular path integration. Nature, 521(7551):186–191, 2015.

[57] Jaison Jiro Omoto, Bao-Chau Minh Nguyen, Pratyush Kandimalla, Jennifer Kelly Lovick, Jeffrey Michael Donlea, and Volker Hartenstein. Neuronal constituents and putative interactions within the *Drosophila* ellipsoid body neuropil. Frontiers in neural circuits, 12:103, 2018.

[58] Tim-Henning Humberg, Pascal Bruegger, Bruno Afonso, Marta Zlatic, James W Truman, Marc Gershow, Aravinthan Samuel, and Simon G Sprecher. Dedicated photoreceptor pathways in drosophila larvae mediate navigation by processing either spatial or temporal cues. Nature communications, 9(1):1260, 2018.

[59] Stanley Heinze. Unraveling the neural basis of insect navigation. Current opinion in insect science, 24:58–67, 2017.

[60] Louis K Scheffer, C Shan Xu, Michal Januszewski, Zhiyuan Lu, Shin-ya Takemura, Kenneth J Hayworth, Gary B Huang, Kazunori Shinomiya, Jeremy Maitlin-Shepard, Stuart Berg, et al. A connectome and analysis of the adult drosophila central brain. elife, 9:e57443, 2020.

[61] Chih-Yung Lin, Chao-Chun Chuang, Tzu-En Hua, Chun-Chao Chen, Barry J Dickson, Ralph J Greenspan, and Ann-Shyn Chiang. A comprehensive wiring diagram of the protocerebral bridge for visual information processing in the drosophila brain. Cell reports, 3(5):1739–1753, 2013.

[62] Corey S Goodman, Michael J Bastiani, Chris Q Doe, Sascha Du Lac, Stephen L Helfand, John Y Kuwada, and John B Thomas. Cell recognition during neuronal development. Science, 225(4668):1271–1279, 1984.

[63] J Roger Jacobs and Corey S Goodman. Embryonic development of axon pathways in the drosophila cns. ii. behavior of pioneer growth cones. Journal of Neuroscience, 9(7):2412–2422, 1989.

[64] Claire Eschbach, Akira Fushiki, Michael Winding, Casey M Schneider-Mizell, Mei Shao, Rebecca Arruda, Katharina Eichler, Javier Valdes-Aleman, Tomoko Ohyama, Andreas S Thum, et al. Recurrent architecture for adaptive regulation of learning in the insect brain. Nature neuroscience, 23(4):544–555, 2020.

[65] Lily Kahsai and Åsa ME Winther. Chemical neuroanatomy of the drosophila central complex: distribution of multiple neuropeptides in relation to neurotransmitters. Journal of Comparative Neurology, 519(2):290–315, 2011.

[66] Lily Kahsai, Mikael A Carlsson, Å ME Winther, and Dick R Nässel. Distribution of metabotropic receptors of serotonin, dopamine, gaba, glutamate, and short neuropeptide f in the central complex of drosophila. Neuroscience, 208:11–26, 2012.

[67] Katrin Vogt, David M Zimmerman, Matthias Schlichting, Luis Hernandez-Nunez, Shanshan Qin, Karen Malacon, Michael Rosbash, Cengiz Pehlevan, Albert Cardona, and Aravinthan DT Samuel. Internal state configures olfactory behavior and early sensory processing in drosophila larvae. Science advances, 7(1):eabd6900, 2021.

[68] Diogo Pimentel, Jeffrey M Donlea, Clifford B Talbot, Seoho M Song, Alexander JF Thurston, and Gero Miesenböck. Operation of a homeostatic sleep switch. Nature, 536(7616):333–337, 2016.

[69] Lucia L Prieto-Godino, Soeren Diegelmann, and Michael Bate. Embryonic origin of olfactory circuitry in drosophila: contact and activity-mediated interactions pattern connectivity in the antennal lobe. 2012.

[70] Arnaldo Carreira-Rosario, Aref Arzan Zarin, Matthew Q Clark, Laurina Manning, Richard D Fetter, Albert Cardona, and Chris Q Doe. Mdn brain descending neurons coordinately activate backward and inhibit forward locomotion. Elife, 7:e38554, 2018.

[71] James W Truman and Lynn M Riddiford. The evolution of insect metamorphosis: a developmental and endocrine view. Philosophical Transactions of the Royal Society B, 374(1783):20190070, 2019.

[72] Jessica Kromp, Tilman Triphan, and Andreas S Thum. Finding a path: Local search behavior of drosophila larvae. bioRxiv, pages 2024–11, 2024.

[73] Roland Strauss and Martin Heisenberg. A higher control center of locomotor behavior in the drosophila brain. Journal of Neuroscience, 13(5):1852–1861, 1993.

[74] Sean M Buchanan, Jamey S Kain, and Benjamin L De Bivort. Neuronal control of locomotor handedness in drosophila. Proceedings of the National Academy of Sciences, 112(21):6700–6705, 2015.

[75] Olga V Alekseyenko, Yick-Bun Chan, Ran Li, and Edward A Kravitz. Single dopaminergic neurons that modulate aggression in drosophila. Proceedings of the National Academy of Sciences, 110(15):6151–6156, 2013.

[76] Jun Tomita, Gosuke Ban, Yoshiaki S Kato, and Kazuhiko Kume. Protocerebral bridge neurons that regulate sleep in drosophila melanogaster. Frontiers in Neuroscience, 15:647117, 2021.

